# Connecting Concepts in the Brain by Mapping Cortical Representations of Semantic Relations

**DOI:** 10.1101/649939

**Authors:** Yizhen Zhang, Kuan Han, Robert Worth, Zhongming Liu

**Affiliations:** Department of Electrical Engineering and Computer Science, University of Michigan, Ann Arbor; Department of Biomedical Engineering, University of Michigan, Ann Arbor; Weldon School of Biomedical Engineering, Purdue University, West Lafayette; School of Electrical and Computer Engineering, Purdue University, West Lafayette; Department of Mathematical Sciences, Indiana University-Purdue University Indianapolis

## Abstract

In the brain, the semantic system is thought to store concepts. However, little is known about how it connects different concepts and infers semantic relations. To address this question, we collected hours of functional magnetic resonance imaging data from human subjects listening to natural stories. We developed a predictive model of the voxel-wise response and further applied it to thousands of new words. Our results suggest that both semantic categories and relations are represented by spatially overlapping cortical patterns, instead of anatomically segregated regions. Semantic relations that reflect conceptual progression from concreteness to abstractness are represented by cortical patterns of activation in the default mode network and deactivation in the frontoparietal attention network. We conclude that the human brain uses distributed networks to encode not only concepts but also relationships between concepts. In particular, the default mode network plays a central role in semantic processing for abstraction of concepts.

## Introduction

Humans can describe the potentially infinite features of the world and communicate with others using a finite number of words. To make this possible, our brains need to encode semantics^1^, infer concepts from experiences^2^, relate one concept to another^3,4^, and learn new concepts^5^. Central to these cognitive functions is the brain’s semantic system^6^. It is spread widely over many regions in the association cortex^7–9^, and it also partially overlaps with the default-mode network^10^. Based on piecemeal evidence from brain imaging studies^11,12^ and patients with focal lesions^13^, individual regions in the semantic system are thought to represent distinct categories or domains of concepts^11,13^ grounded in perception, action, and emotion systems^14,15^.

However, little is known about how the brain connects concepts and infers semantic relations^16,17^. As concepts are related to one another in the real world, cortical regions that represent concepts are also connected, allowing them to communicate and work together as networks^18^. It is thus likely that the brain represents semantic relations as emerging patterns of network interaction^19^. Moreover, since different types of concepts may express similar relations, it is also possible that the cortical representation of a semantic relation may transcend any specific conceptual domain. Testing these hypotheses requires a comprehensive study of the semantic system as a set of distributed networks, as opposed to a set of isolated regions. Being comprehensive, the study should also survey cortical responses to a sufficiently large number of words from a wide variety of conceptual domains^1^, ideally using naturalistic stimuli^20^.

Similar to a prior work^1^, we developed a predictive model of human functional magnetic resonance imaging (fMRI) responses given >11 hours of natural story stimuli. In this model, individual words and their pairwise relationships were both represented as vectors in a continuous semantic space^21^, which was learned from a large corpus and was linearly mapped onto the brain’s semantic system. Applying this model to thousands of words and hundreds of word pairs, we have demonstrated the distributed cortical representations of semantic categories and semantic relations, respectively. Our results also shed new light on the role of the default mode network in semantic processing.

## Results

### Word embeddings predicted cortical responses to speech

To extract semantic features from words, we used a word2vec model trained to predict the nearby words of every word in large corpora^21^. Through word2vec, we could represent any word as a vector in a 300-dimensional semantic space. Of this vector representation (or word embedding), every dimension encoded a distinct semantic feature learned entirely by data-driven methods^21^, instead of by human intuition or linguistic rules^1,22,23^. To relate this semantic space to its cortical representation, we defined a voxel-wise encoding model^24^ – a multiple linear regression model that expressed each voxel’s response as a weighted sum of semantic features^1^ (Figure 1).

**Figure 1.**
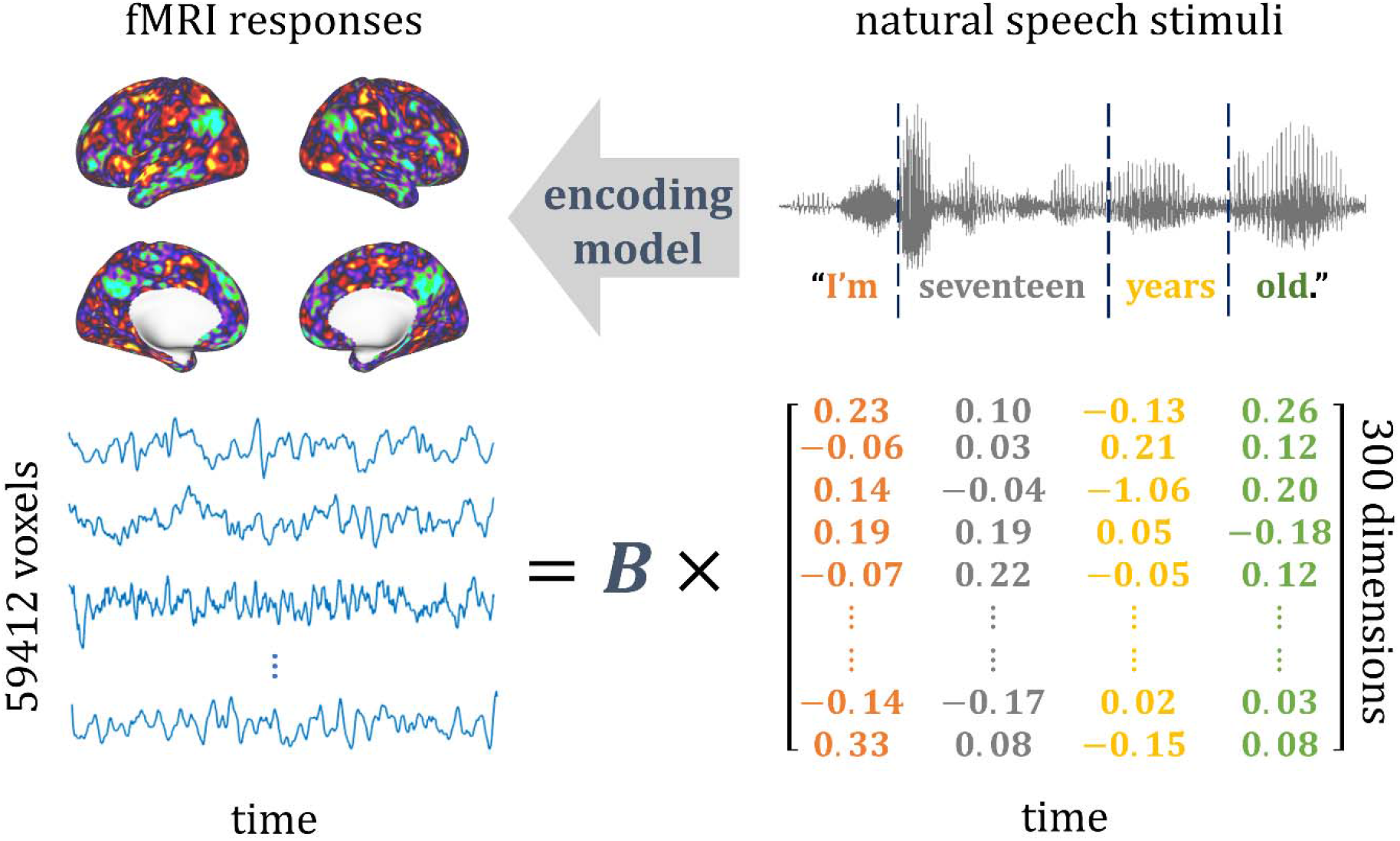
Illustration of the encoding model. The encoding model was trained and tested for predicting the fMRI responses (top-left) to a time series of words in audio story stimuli (top-right). Every word (as color coded) was converted to a 300-dimensional vector through word2vec. The encoding model was denoted as a 59,421-by-300 matrix (B) to predict the voxel response to every word (bottom).

To estimate the voxel-wise encoding model, we acquired whole-brain fMRI data from 19 native English speakers listening to different audio stories (from The Moth Radio Hour: https://themoth.org/radio-hour), each repeated twice for the same subject. By counterbalancing the stories across subjects, we sampled different words with different subjects, such that the sampled words for every subject covered similar distributions in the semantic space (Supplementary Figure 1) and included a common set of frequent words (Supplementary Figure 2 and Supplementary Table 1), while every semantic category or relation of interest was sampled roughly evenly across subjects (Supplementary Figure 3 & 4). In total, the story stimuli combined across subjects lasted 11 hours and included 47,356 words (or 5,228 words if duplicates were excluded). The voxel-wise encoding model was estimated based on the fMRI data concatenated across all stories and subjects.

By 10-fold cross-validation^25^, the model-predicted response was significantly correlated with the measured fMRI response (block-wise permutation test, false discovery rate or FDR *q* < 0.05) for voxels broadly distributed on the cortex (Figure 2). The voxels highlighted in Figure 2 were used to delineate an inclusive map of the brain’s semantic system, because the cross-validation was applied to a large set of (5,228) words, including those most frequently used in daily life (Supplementary Figure 2). This map, hereafter referred to as the semantic system, was widespread across regions from both hemispheres, as opposed to only the left hemisphere, which has conventionally been thought to dominate language processing and comprehension^26^.

**Figure 2.**
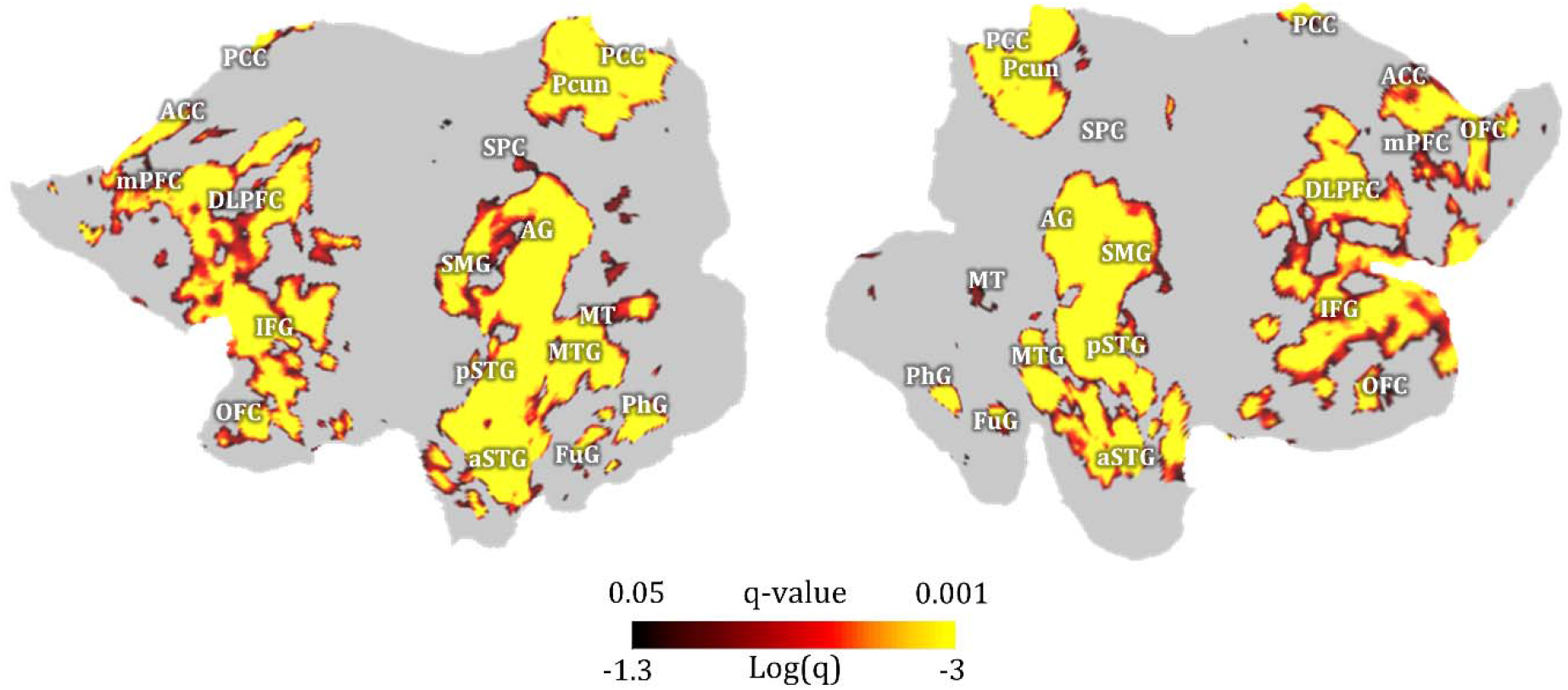
A map of the semantic system obtained by 10-fold cross-validation of the encoding model. The map is displayed on the flattened cortical surfaces for the left and right hemispheres. The color indiciates the FDR (or *q*-value) in a logarithmic scale. The highlighted areas include voxels where crossvalidation results are statistically significant (block-wise permutation test, one-sided, *q* < 0.05). Anatomical labels are shown to relate the model-predictable areas to cortical regions predefined in a connectivity-based atlas^27^.

We also tested how well the trained encoding model could be generalized to a new story never used for model training and whether it could be used to account for the differential responses at individual regions. For this purpose, we acquired the voxel response to an independent testing story (6 mins 53 secs, 368 unique words) for every subject and averaged the response across subjects. As shown in Figure 3a, we found that the encoding model was able to reliably predict the evoked responses in the inferior frontal sulcus (IFS), supramarginal gyrus (SMG), angular gyrus (AG), superior temporal gyrus (STG), middle temporal visual area (MT), left fusiform gyrus (FuG), left parahippocampal gyrus (PhG), and posterior cingulate cortex (PCC) (block-wise permutation test, FDR *q* < 0.05). These regions of interest (ROIs), as predefined in the human brainnetome atlas^27^ (Figure 3b, see details in Supplementary Table 2), showed different response dynamics given the same story, suggesting their highly distinctive roles in semantic processing (Figure 3c). Despite such differences across regions, the encoding model was found to successfully predict the response time series averaged within every ROI except the right FuG (Figure 3c), suggesting its ability to explain the differential semantic coding (i.e. stimulus-response relationship) at different regions.

**Figure 3.**
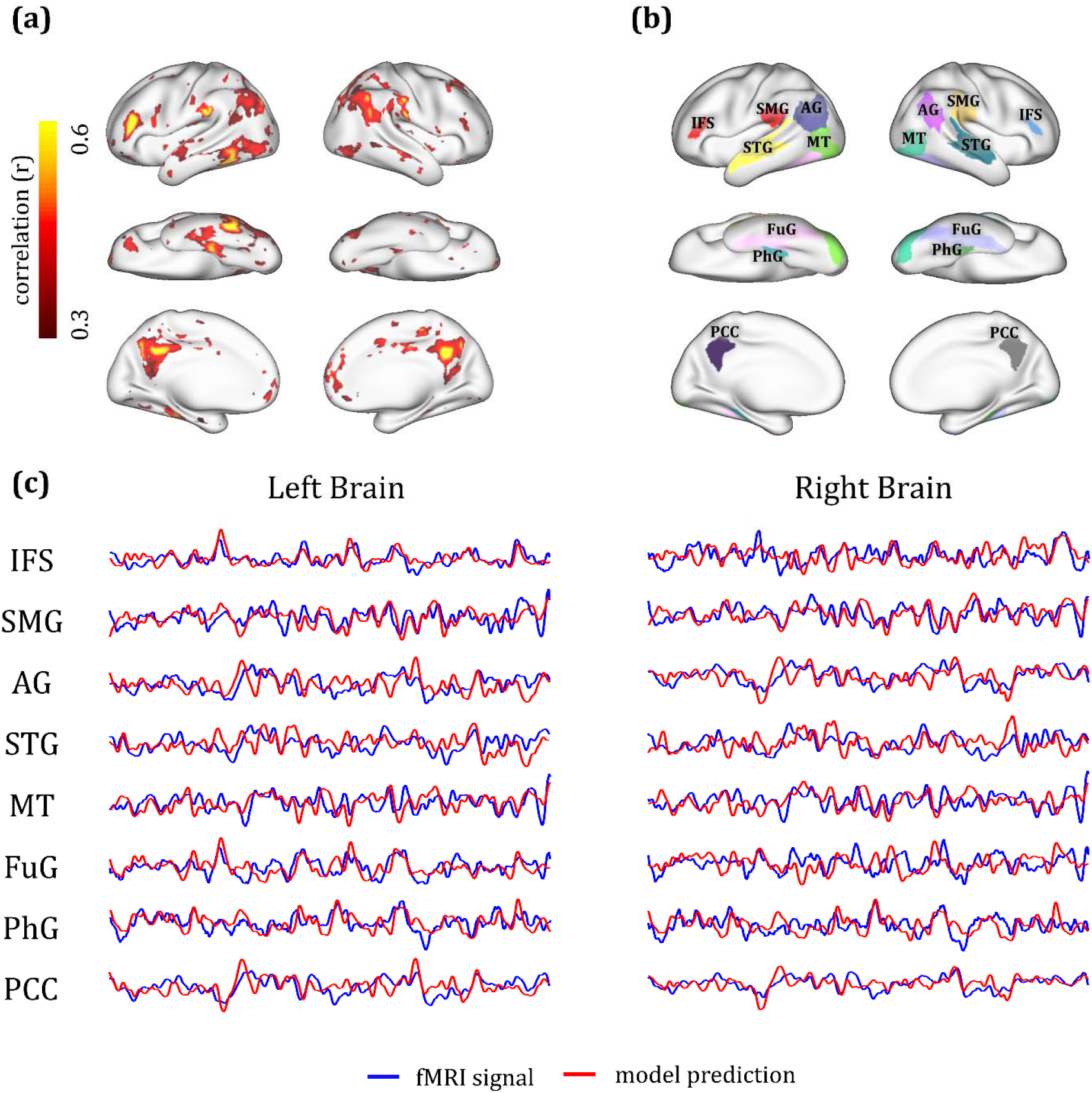
Measured vs. model-predicted responses to a new (untrained) testing story. (a) The voxelwise correlation between fMRI responses and model predictions for a 6min 53sec testing story. The fMRI responses were averaged across subjects. Color indicates the correlation coefficient. The color-highlighted areas include the voxels of statistical significance (block-wise permutation test, one-sided, FDR *q* < 0.05). (b) Pre-defined ROIs (shown in different colors) are displayed on the cortical surfaces. See details in Supplementary Table 2. (c) Response time series as measured (blue) or model-predicted (red) for each ROI, by averaging the time series across voxels within each ROI. IFS, inferior frontal sulcus; SMG, supramarginal gyrus; AG, angular gyrus; STG, superior temporal gyrus; MT, middle temporal visual area; FuG, fusiform gyrus; PhG: parahippocampal gyrus; PCC, posterior cingulate gyrus.

### Distributed cortical patterns encoded semantic categories

Since the encoding model was generalizable to new words and sentences, we further used it to predict cortical responses to >9,000 words from nine categories: *tool, human,plant, animal,place, communication, emotion, change, quantity* (Supplementary Table 3), as defined in WordNet^28^ and are representative of different conceptual domains. We confined the model prediction to the voxels in the semantic system for which the model fit was significant during cross-validation (Figure 2). Within each category, we averaged the model-predicted responses given every word and mapped the statistically significant voxels (one sample t-test, FDR *p* < 0.01). This map represented each category being projected from the semantic space to the cortex, and thus was interpreted as the model-predicted cortical representation of each category. We found that individual categories were represented by spatially overlapping and distributed cortical patterns (Figure 4). For example, the category *tool* was represented by the SMG, posterior middle temporal gyrus (pMTG), FuG, and inferior frontal gyrus (IFG); this representation was more pronounced in the left hemisphere than the right hemisphere. Such categories as *human, plant*, and *animal* were also represented more by the left hemisphere than the right hemisphere. The category *place* was represented by bilateral PhG, dorsolateral prefrontal cortex (dLPFC), and AG. In contrast, *communication, emotion, change*, and *quantity*, showed stronger representations in the right hemisphere than in the left hemisphere. Although the size of word samples varied across categories (Supplementary Table 3), the sample size was sufficiently large for every category, since the resulting category representation had reached or approached its maximum extent at the given sample size (Supplementary Figure 5). See Supplementary Method 6 for more details about testing the effect of sample size on categorical representation.

**Figure 4.**
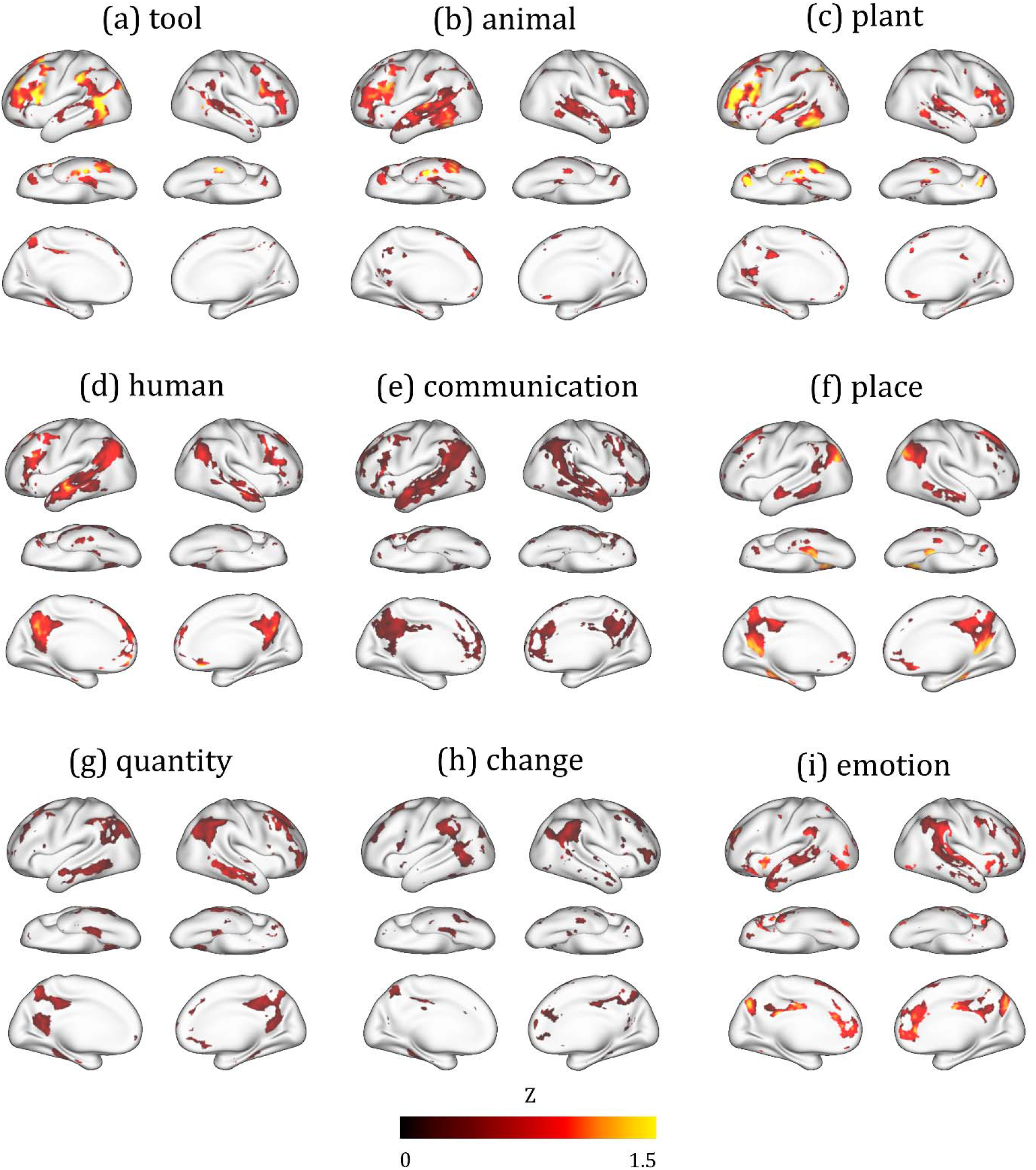
Cortical representations of semantic categories. For each category, the color indicates the mean of the normalized response (or the z score) averaged across word samples in the category (Supplementary Table 3). The color-highlighted areas include the voxels of statistical significance (one sample t-test, one-sided, FDR *q* < 0.01).

To each voxel in the semantic system, we assigned a single category that gave rise to the strongest voxel response, thus dividing the semantic system into category-labeled parcels (Figure 5a). The resulting parcellation revealed how every category of interest was represented by a different set of regions, as opposed to any single region. In addition, the distinction in left/right lateralization was noticeable and likely attributable to the varying degree of concreteness for the words from individual categories. The concepts lateralized to the left hemisphere appeared relatively more concrete or exteroceptive, whereas those lateralized to the right hemisphere were more abstract or interoceptive (Figure 5b). This intuitive interpretation was supported by human rating of concreteness (from 1 to 5) for every word in each category^29^. The concreteness rating was high (between 4 and 5) for the categories lateralized to the left hemisphere, whereas it tended to be lower yet more variable for those categories dominated by the right hemisphere (Figure 5c).

**Figure 5.**
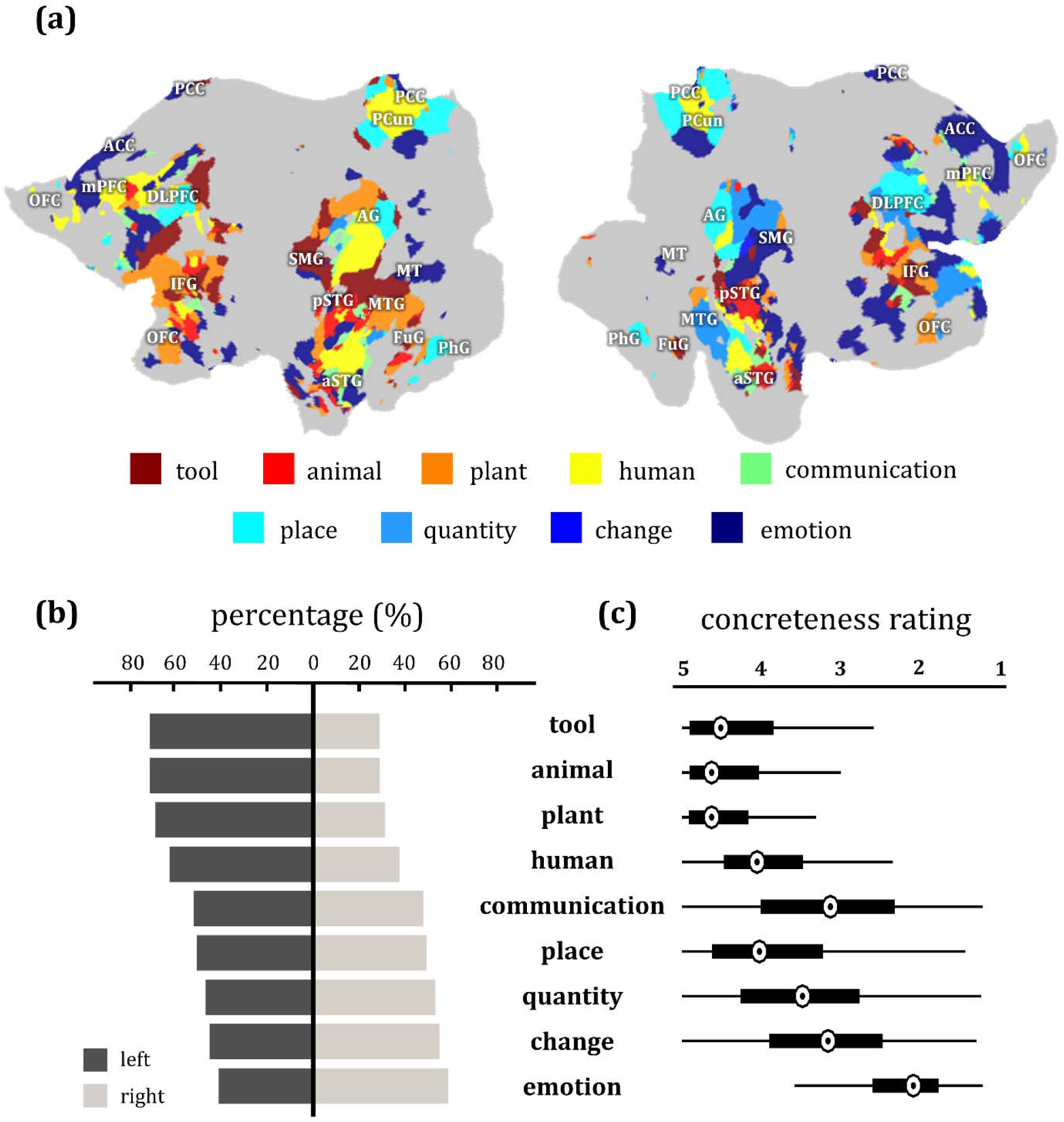
Cortical representation of semantic category. (a) Category-labeled parcellation based on voxel-wise selectivity using a “winners-take-all” strategy. (b) Cortical lateralization of categorical representations. For each category, the percentage value was calculated by counting the number of voxels on each hemisphere that represented the given category. The number of word samples in each of the nine categories is: tool (200), animal (734), plant (387), human (808), communication (2027), place (814), quantity (958), change (3417), and emotion (504). (c) The concreteness rating of words in each category. The maximum value of the concreteness rating is 5.00 and the minima value is 1.25. In this box plot, the central mark indicates the median, and the box edges indicate the 25th and 75th percentiles respectively. The maximum whisker length is 1.5.

### Co-occurring activation & deactivation encoded word relation

Through the word2vec model, we could also represent semantic relations as vectors in the semantic space^30^. Specifically, we represented the relationship between any pair of words based on their difference vector in word embedding. We chose word-pairs from the SemEval-2012 Task 2 dataset^31^. Every chosen word-pair had been human rated as an affirmative example of one of 10 classes of semantic relation: *whole-part, class-inclusion, object-attribute, case relations, space-associated, time-associated, similar, contrast, object-nonattribute*, and *cause-effect* (Supplementary Table 4). For the first 6 classes, the relation vectors in the semantic space were found to be more consistent across word-pairs in the same class than those in different classes (Supplementary Figure 6). For each of the first 6 classes, the relation between every pair of words could be better identified based on the relation vectors of the other word-pairs in the same class than those from any different class, showing the top-1 identification accuracy from 40% to 83% against the 10% chance level (Supplementary Figure 7).

For a given word-pair, their relation vector could be further projected onto the cortex through the encoding model. For an initial exploration, we applied this analysis to 178 word-pairs that all shared a same *whole-part* relationship. For example, in four word-pairs, (hand, finger), (zoo, animal), (hour, second), and (bouquet, flower), *finger* is part of *hand; animal* is part of *zoo*; *second* is part of *hour; flower* is part of *bouquet*. Individually, the words from different pairs had different meanings and belonged to different semantic categories, as *finger, animal, second*, and *flower* were semantically irrelevant to one another. Nevertheless, their pairwise relations all entailed the *whole-part* relation, as illustrated in Figure 6a. By using the encoding model, we mapped the pairwise word relationship onto voxels in the semantic system (as shown in Figure 2), averaged the results across pairs, and highlighted the significant voxels (paired permutation test, FDR *q* < 0.05). The resulting cortical map represented each semantic relation being projected from the semantic space to the cortex, reporting the model-predicted cortical representation of the relation. We found that the *whole-part* relation was represented by a cortical pattern that manifested itself as the co-occurring activation of the default mode networks^32^ (DMN, including AG, MTG, and PCC) and deactivation of the frontoparietal network^33,34^ (FPN, including LPFC, IPC and pMTG) (Figure 6b). This cortical pattern encoded the *whole-part* relation independent of the cortical representations of the individual words in this relation. The co-activation and deactivation pattern indicated that conceptual progression from *part* to *whole* manifested itself as increasing deactivation of FPN alongside increasing activation of DMN, whereas progression from *whole* to *part* was shown as the reverse cortical pattern varying in the opposite direction, as illustrated in Figure 6c.

**Figure 6.**
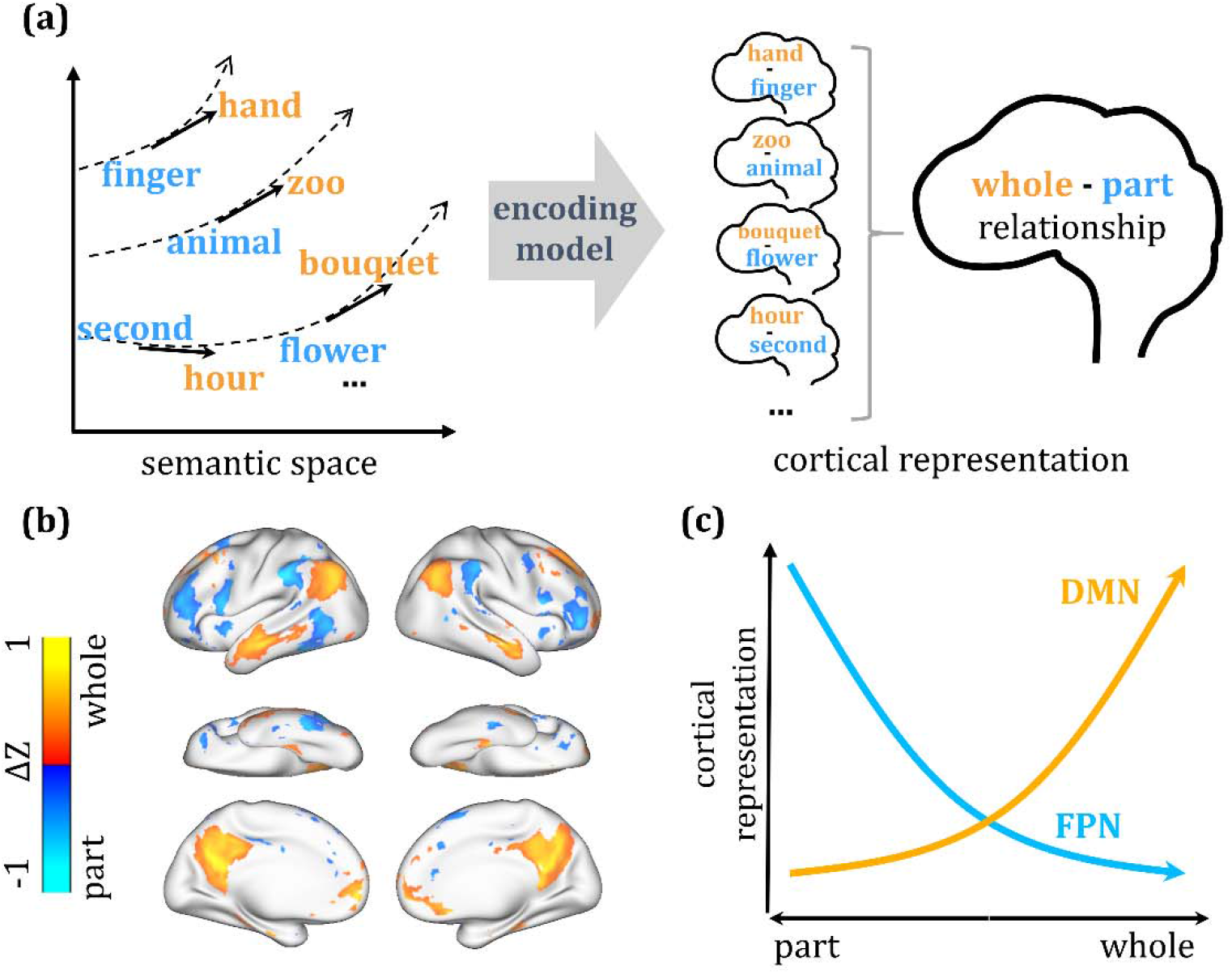
Mapping cortical representation of the whole-part relation. (a) The illustration of mapping the whole-part relation from the semantic space to the human brain through the voxel-wise encoding model. We viewed the whole-part relation as a vector field over the semantic space. This relation field was sampled by the difference vector of each word pair that held such a relation (left). The cortical representation of this difference vector was predicted by the voxel-wise encoding model. Cortical representation of the whole-part relation was then obtained by averaging representations of all word pairs (right). (b) Cortical representation of the whole-part relation. The statistical significance was assessed by a paired permutation test (178 word pairs, two-sided, FDR *q* < 0.05). (c) The co-occurring activation of DMN and deactivation of FPN encodes the whole-part relation or the conceptual progression from part to whole.

Similarly, we also mapped the cortical representations of several other semantic relations. Each relation was projected to a distinct cortical pattern (Figure 7). Specifically, the *class-inclusion* relation, e.g. (color, green) where *color* includes *green*, was represented by the activation of AG and MTG and the deactivation of IFG and STG (Figure 7b). The *object-attribute* relation, e.g. (fire, hot) where *fire* is *hot*, was represented by an asymmetric cortical pattern including activation primarily in the left hemisphere and deactivation primarily in the right hemisphere (Figure 7c). The *case relations*, e.g. (coach, player) where a *coach* teaches a *player*, was represented by a cortical pattern similar to that of the *whole-part* relation (Figure 7d), despite a lack of intuitive connection between the two relations. The *space-associated* relation, e.g. (library, book) where *book* is an associated item in a *library*, was represented by activation of AG and PCC and deactivation of STG (Figure 7e). Lastly, the *time-associated* relation, e.g. (morning, sunrise) where *sunrise* is a phenomenon associated with *morning*, was also represented by a bilaterally asymmetric pattern (Figure 7f). A graph-based illustration of the representational geometry further highlights the distinction across semantic relations in terms of their bilateral (a)symmetry and engagement of individual ROIs (Supplementary Figure 8). However, several nominal (human-defined) relations, e.g. *similar, contrast, object-nonattribute* and *cause-effect*, were projected onto either no or fewer voxels (Supplementary Table 4 & Supplementary Figure 9).

**Figure 7.**
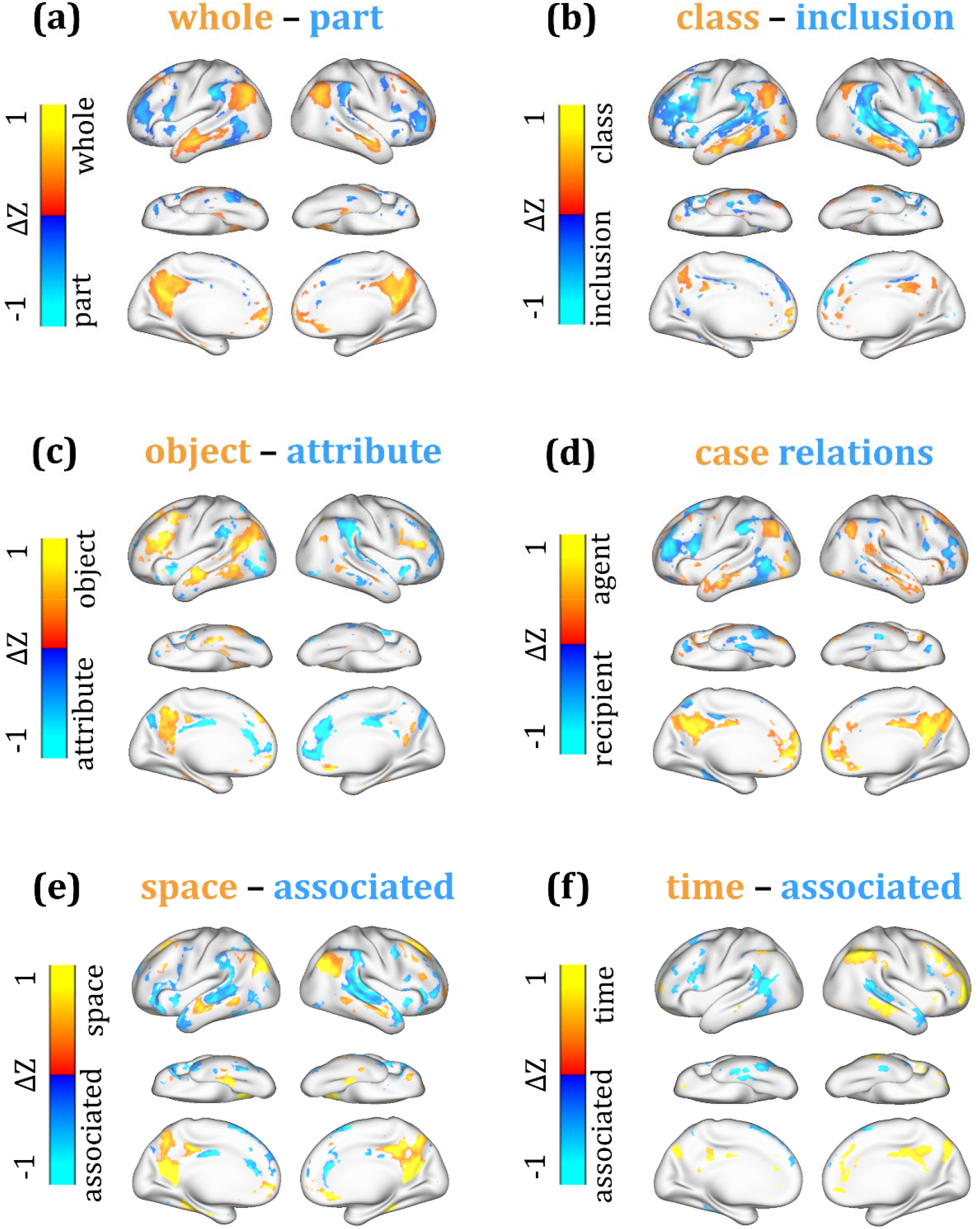
Cortical representations of semantic relations. The cortical pattern associated with each relation shows the average cortical projection of every word-pair sample in that relation and highlights only the voxels of statistical significance (paired permutation test, two-sided, FDR *q* < 0.05) based on voxel-wise univariate analysis. For each of the six relations in this figure, see Supplementary Table 4 for summary statistics and also see Supplementary Figure 9 for the results about other relations.

The voxel-wise univariate analysis restricted the representation of each semantic relation to one cortical pattern while ignoring the interactions across voxels and regions. This limitation led us to use a principal component analysis to decompose the cortical projection of the difference vector of every word pair in each semantic relation. This multivariate analysis revealed two cortical patterns that were statistically significant (one-sample t-test, *p* < 0.01) in representing the semantic relation of *object-attribute, case relations*, or *space-associated*, but only revealed one pattern for the relation of *whole-part, class-inclusion, time-associate*, or *cause-effects* (Supplementary Figure 9). Interestingly, when two cortical patterns represented the same semantic relation, they seemed to correspond to different sub-classes of that relation (Figure 8). For the relation of *object-attribute*, one cortical pattern corresponded to *inanimate object-attribute*, e.g. (candy, sweet), and the other corresponded to *human-attribute*, e.g. (coward, fear) (Figure 8a). Similarly, the two cortical patterns for *case relations* corresponded to *agent-instrument* and *action-recipient*, respectively (Figure 8b). The *space-associated* relation was distinctively represented for its two sub-classes: *space-associated item* and *space-associated activity* (Figure 8c). The cortical patterns that represented a relation, as obtained with either the multivariate or univariate analysis, highlighted generally similar regions (Supplementary Figure 9).

**Figure 8.**
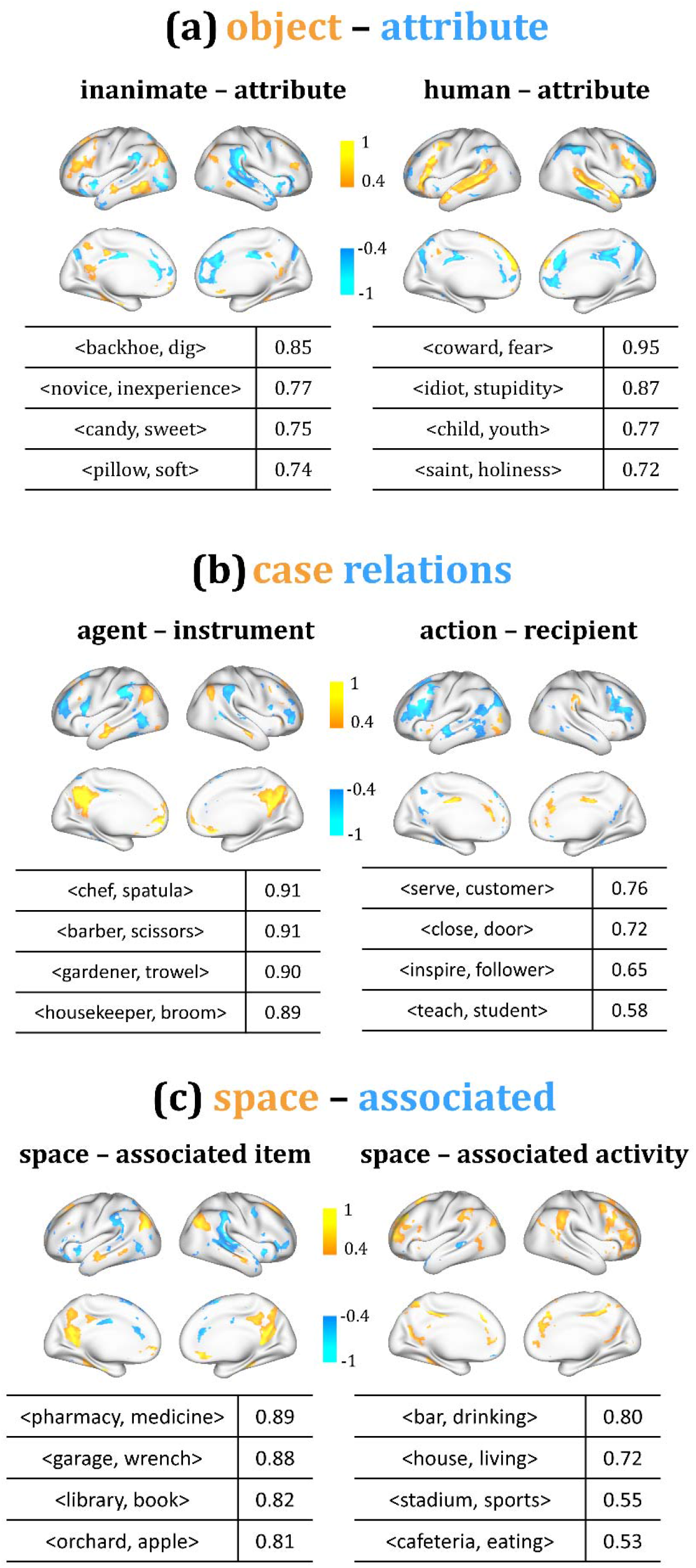
Multiple cortical patterns represent sub-classes of individual semantic relations. For three semantic relations: “object-attribute” (a), “case relations” (b) and “space-associated” (c), multivariate pattern analysis revealed two cortical patterns associated with each of these relations. The cortical patterns were min-max normalized to [-1, 1]. The table lists the top-4 word-pairs, of which the cortical projection was most similar (in terms of cosine-similarity) with the first (left) or second (right) cortical pattern associated with each relation. See Supplementary Figure 9 for results about other relations.

## Discussion

Using fMRI data from subjects listening to natural story stimuli, we established a predictive model to map the cortical representations of semantic categories and relations. We found that semantic categories were not represented by segregated cortical regions but instead by distributed and overlapping cortical patterns, mostly involving multimodal association areas. Although both cerebral hemispheres supported semantic representations, the left hemisphere was more selective to concrete concepts whereas the right hemisphere was more selective to abstract concepts. Importantly, semantic relations were represented by co-occurring activation and deactivation of distinct cortical networks. Semantic relations that reflected conceptual progression from concreteness to abstractness were represented by the co-occurrence of activation in the default-mode network and deactivation in the attention network. Interestingly, some semantic relations could each be represented by two cortical patterns, corresponding to intuitively distinct sub-classes of the relation. Our findings suggest that the human brain represents a continuous semantic space. To support conceptual inference and reasoning, the brain uses distributed cortical networks to encode not only concepts but also relationships between concepts. Notably, the default-mode network plays an active role in semantic processing for abstraction of concepts. In the following, we discuss our methods and findings from the joint perspective of machine learning and neuroscience in the context of natural language processing.

Central to this study is the notion of embedding concepts in a continuous semantic space^21^. Although we use words to study concepts, words and concepts are different. The vocabulary is finite, but concepts are infinite. We view words vs. concepts as discrete vs. continuous samples from the semantic space. Moving a concept in the semantic space may create a new concept or arrive at a different concept. This provides the flexibility for using concepts to describe the world, as is understood by the brain and, to a lesser extent, as is expressed in language.

Moreover, concepts are not isolated but related to one another. Since we view concepts as points in the semantic space, we consider conceptual relationships to be continuous vector fields in the same space. A position in the semantic space may experience multiple fields, and different positions may experience the same field. Thus, a concept may relate to other concepts in various ways, and different pairs of concepts may hold the same relation^4^. Because semantics reflect how humans understand and describe the world, we hypothesize that the brain not only encodes such a continuous semantic space^1^ but also encodes semantic relations as vector fields in the semantic space.

Machine learning leverages the notion of continuous semantic space for natural language processing^21,35,36^, and provides a new way to model and reconstruct neural responses^1,24,37–39^. For example, word2vec can represent millions of words as vectors in a lower-dimensional semantic space^21^. Two aspects of word2vec have motivated us to use it for this study. First, words with similar meanings share similar linguistic contexts and have similar vector representations in the semantic space^40^. Second, the relationship between two words is represented by their difference vector, which is transferable to another word. For an illustrative example, “(*man - women*) + *queen*” results in a vector close to “*king*”^30^. As such, word2vec defines a continuous semantic space and preserves both word meanings and word-to-word relationships.

In addition, word2vec learns the semantic space from large corpora in a data-driven manner^21^. This is different from defining the semantic space based on keywords that are hand selected^22^, frequently used^41^, minimally grounded, or neurobiologically relevant^23,42^. Although those word models are seemingly more intuitive, they are arguably subjective and may not be able to describe the complete semantic space. We prefer word2vec as a model of word embedding, because it leverages big data to learn natural language statistics without any human bias. We assume that the brain encodes a continuous semantic space similarly as is obtained by word2vec. Since word2vec is not constrained by any neurobiological knowledge, we do not expect it to encode the exactly same semantic space as does the brain. Instead, we hypothesize that the word2vec-based semantic space and the brain are similar up to linear projection (i.e. transformation through linear encoding).

Our results support this hypothesis and reveal a distributed semantic system (Figure 2). In this study, the semantic system mapped with natural stories and thousands of words resembles the semantic system mapped with meta-analysis of the activation foci associated with fewer words^6^. As in that paper, our results also highlight a similar set of semantics-encoded regions (Figure 2), most of which are associated with high-level integrative processes that transcend any single modality^8,9^. However, our map of the semantic system is largely bilateral, rather than being dominated by the left hemisphere as suggested by Binder et al.^6^, although the activation foci analyzed by Binder et al. are actually distributed on both hemispheres (see Figure 2 in Binder et al.^6^). Importantly, the two hemispheres seem to be selective to different aspects of semantics. Unlike prior findings^43,44^, our results suggest that the left hemisphere tends to encode exteroceptive and concrete concepts, whereas the right hemisphere tends to encode interoceptive and abstract concepts (Figure 4 & 5).

Our semantic system shows a cortical pattern similar to that reported by Huth et al.^1^. This similarity is not surprising, because both studies use similar natural speech stimuli and encoding models. However, unlike Huth et al.^1^, we do not emphasize the semantic selectivity of each region or tile the cortex into regions associated with distinct conceptual domains. On the contrary, none of the conceptual categories addressed in this study is represented by a single cortical region. Instead, individual categories are represented by spatially distributed and partly overlapping cortical networks (Figure 4), each of which presumably integrates various domain-defining attributes by connecting the regions that encode different attributes^11,14,15^. In this regard, our results lend support to efforts that address semantic selectivity by means of networks, as opposed to regions^18,19^.

The primary focus of this study is on semantic relations between words. Extending the earlier discussion about the semantic space, the relationship between words is represented by their vector difference, of which the direction and magnitude indicate different aspects of the relationship. Let us use (minute, day) as an example. Of their relation vector, the direction indicates a part-to-whole relation, and the magnitude indicates the offset along this direction. Starting from *minute* and relative to *day*, a larger offset leads toward *month* or *year*, a smaller offset leads toward *hour*, and a negative offset leads toward *second*. Our results suggest that the direction of relation vector tends to be generalizable and transferrable across word pairs in the same semantic relation (Supplementary Figure 6 & 7). This leads us to hypothesize that the semantic space includes continuous vector fields, each of which represents a semantic relation and is likely applicable to various concepts or even domains of concepts. When a vector field is visualized as many field lines, the points (i.e. concepts) that each field line passes through are related to one another by the same semantic relation (as illustrated by Figure 6a).

In a nominal relation (e.g. *whole-part)*, each word pair takes a discrete sample from the underlying vector field (Figure 6a). Projecting a number of such relational samples onto the cortex reveals one or multiple cortical patterns that encode the relation. Such cortical patterns often manifest themselves as co-occurring activation and deactivation of different regions (Figure 6, 7 & 8). We interpret this co-activation and co-deactivation as an emerging pattern when the brain relates two concepts that hold a meaningful relation, reflecting the progression from one concept to the other. This pattern encodes generalizable differential relations between concepts, as opposed to concepts themselves, serving differential coding that transcends any conceptual domain or category (Figure 6). Speculatively, this network-based coding of semantic relation is an important mechanism that supports analogical reasoning^45^, e.g. matching similar relations with different word-pair samples^31^. This plausible mechanism of the brain might further facilitate humans learning new concepts by connecting them to existing concepts through established semantic relations.

It is also noteworthy that a semantic relation as defined by human intuition may not exactly match the relation as represented by the brain. It is possible that a nominal relation may be heterogeneous and contain multiple sub-classes each being represented by a distinct cortical pattern. Results obtained with multivariate pattern analysis support this notion (Figure 8). It is also reasonable that a nonsensical relation, e.g. *object-nonattribute*, does not have any cortical representation (Supplementary Figure 9).

Although we view word relations and categories as distinct aspects of the semantic space, the two aspects may engage similar cortical networks under specific circumstances. For example, our results indicate that the *space-associated* relation and the *place* category are represented by similar cortical patterns (Figure 4f & Figure 7e). This is unsurprising because the *space-associated* word pairs are often associated with *place*. Such a relation-category association is intrinsic to natural language statistics, and similarly applies to *time-associated* and *quantity*. This does not imply that the semantic relations are always associated with specific semantic categories (Supplementary Figure 10). There is no evidence for a generalizable relation-category association. See Supplementary Method 5 for more details about evaluating the association between relations and categories.

We interpret the co-activation/deactivation patterns as “anti-correlated networks” with respect to the cortical representations of semantic relations. This interpretation is reasonable given the notion of “activate together wire together”. Task-related patterns of cortical activation resemble those emerging from spontaneous activity or resting state networks^46^. In the context of semantics, the anti-correlated networks reported herein encode a semantic relation, or the direction in which one concept relates to another. For example, conceptual progression from *part* to *whole* has a cortical signature as co-occurring activation in DMN and deactivation in FPN (Figure 6b). The opposite direction from *whole* to *part* involves the same regions or networks but reverses their polarity in terms of activation or deactivation. In this example, the cortical co-activation/deactivation pattern is nearly identical to the anti-correlated networks observed with resting state fMRI^47^, and therefore it is likely to be intrinsic and supported by underlying structural connections.

Our results suggest that DMN is involved in cortical processing of not only concepts but also semantic relations. This finding underscores the fact that DMN plays an active role in language and cognition^10,48–51^, rather than only a task-negative and default mode of brain function^32^. In particular, several semantic relations, such as *whole-part* and *class-inclusion*, are all mapped onto DMN (Figure 6 & 7), suggesting that DMN is likely associated with the semantic regularity common to those relations. Indeed, words being *whole* or *class* are more abstract and general, whereas those being *part* or *inclusion* are relatively more detailed and specific. As such, these relations all indicate (to a varying degree) conceptual abstraction. This progression involves DMN, increasing or decreasing its activity as a concept (of various types) becomes more abstract or specific, respectively. Moreover, concrete concepts, e.g. *tool, plant, animal*, are represented by regions outside the DMN, whereas more abstract concepts, e.g. *communication, emotion, quantity*, are represented by cortical regions that reside in, or at least overlap with, DMN (Figure 4 & 5).

These observations lead us to speculate that DMN underlies a cognitive process for abstraction of concepts. This interpretation is consistent with findings from several prior studies^51,52^. For example, Spunt et al. has shown that conceptualizing the same action at an increasingly higher level of abstraction gives rise to an increasingly greater responses at regions within DMN^52^. Sormaz et al. has shown evidence that activity patterns in DMN during cognitive tasks are associated with whether thoughts are detailed, rather than whether they are task related or unrelated^51^. In contrast to DMN, another network, FPN, seems to play an opposite role in semantic processing. FPN is often activated by attention-demanding tasks and is intrinsically anti-correlated with DMN^47^. Our results suggest that FPN is increasingly activated when the brain is engaged in conceptual specification.

Although our experimental design is justifiable by practical and methodological considerations, it is worth further noting potential limitations, additional justifications and future directions. In this study, we used different stories for different subjects to collect a large set (47,356 words) of stimulus-response samples for training the encoding model. It is logistically difficult to acquire enough data from a single subject. A typical fMRI experiment lasts <2 hours to avoid fatigue, while >11 hours of fMRI scans as needed for the desired sample size would be too long to be realistic. This study design might be of potential concern that individual differences, e.g. laterality^26,53^, are confounded with the words used for model training. If a subset of subjects is over-represented for one semantic dimension and a different subset is over-represented for a different dimension, the trained encoding model would reflect the idiosyncratic variation across individuals. To mitigate this concern, we had counterbalanced the stories across subjects. By counterbalancing, the stories for different subjects similarly sampled the semantic space (Supplementary Figure 1), the semantic categories or relations of interest (Supplementary Figure 3 & 4), as well as a common set of frequently used words (Supplementary Figure 2). In addition, the use of audio stories as naturalistic stimuli gave rise to highly reproducible cortical responses across subjects, as shown in prior studies^20^ and reinforced by our results (Supplementary Figure 11). See Supplementary Method 2 for more details on testing effects of individual variance.

It might appear counterintuitive that some intuitive semantic relations, e.g. *similar* and *contrast*, did not map onto any informative voxels despite an adequate sample size (Supplementary Figure 9). In fact, it is not surprising at all because such relations are both symmetric. For example, (hot, cold) holds a *contrast* relation, while (cold, hot) also holds the same relation. Likewise, the *similar* relation is also symmetric. In contrast, other relations, e.g. *whole-part*, and *case-relations*, are asymmetric. The relation is directed such that flipping two words in a pair changes the relation. Since we use differential vectors to evaluate word relations, our method is more suited for addressing asymmetric relations, instead of symmetric relations.

In this study, the sample size varied across categories or relations. A potential concern might be that the varying sample size could influence the area to which a category or relation was projected. However, this was not a flaw in study design and did not invalidate our findings. Note that the sample-size difference is intrinsic to how English words are distributed across categories or relations. It was our intention to limit our samples to established datasets from published studies with human behavioral data available and associated with words or word relations^29,31^. Moreover, there was no significant correlation between the sample size and the number of voxels that represented a category (*r* = 0.0017, *p* = 0.99) or relation (*r* = –0.24; *p* = 0.50).

Central to this study, we bridge linguistic models and fMRI data during naturalistic audio-story stimuli. Our findings about cortical representations of semantic categories or relations are based on generalizing a predictive model beyond the data used to train the model. While the generalization is supported by our results on model cross-validation and testing, it is desirable to validate some of our model-predicted findings with experimental data in future studies. Importantly, our computational model-based strategy enables high-throughput investigation of how the brain encodes concepts and relations beyond what is feasible for a single experiment. Hypotheses informed by the model may also lend inspiration to future experimental studies. Moreover, it will also be useful to incorporate neurobiological principles and refines the word2vec model in order to improve the correspondence between the word embedding and the word representation on the human cortex.

## Methods

### Subjects, stimuli and experiments

19 human subjects (11 females, age 24.4±4.8, all right-handed) participated in this study. All subjects provided informed written consent according to a research protocol approved by the Institutional Review Board at Purdue University. While being scanned for fMRI, each subject was listening to several audio stories collected from The Moth Radio Hour (https://themoth.org/radio-hour) and presented through binaural MR-compatible headphones (Silent Scan Audio Systems, Avotec, Stuart, FL). A single story was presented in each fMRI session (6 mins 48 secs ± 1 min 58 secs). For each story, two repeated sessions were performed for the same subject.

Different audio stories were used for training vs. testing the encoding model. For training, individual subjects listened to different sets of stories. When combined across subjects, the stories used for training amounted to a total of 5 hours 33 mins (repeated twice). This design provided a large number of stimulus-response samples beneficial for training the encoding model, which aimed to map hundreds of semantic features to thousands of cortical voxels. For testing, every subject listened to the same single story for 6 mins 53 secs; this story was different from those used for training.

In an attempt to sample a sufficiently large number of words in the semantic space, we intentionally chose audio stories of diverse contents. Since different subjects listened to distinct (training) stories, we further counterbalanced the stories across subjects. For different subjects, the stories included different words (2,492 ± 423) but sampled similar distributions in the semantic space (Supplementary Figure 1)^54^. For each semantic category or relation of interest, the associated words were roughly evenly sampled across subjects (Supplementary Figure 3 & 4). The stories presented to each subject also included a set of common words used frequently in daily life (Supplementary Figure 2). In total, the training stories include 5,228 unique words. By counterbalancing the stories across subjects, we attempted to avoid any notable sampling bias that could significantly confound the idiosyncratic variation across subjects with the variation of the sampled words across subjects. See Supplementary Method 1 for more details.

### Data acquisition and processing

T_1_ and T_2_-weighted MRI and fMRI data were acquired in a 3T MRI system (Siemens, Magnetom Prisma, Germany) with a 64-channel receive-only phased-array head/neck coil. The fMRI data were acquired with 2 mm isotropic spatial resolution and 0.72 s temporal resolution by using a gradient-recalled echo-planar imaging sequence (multiband = 8, 72 interleaved axial slices, TR = 720 ms, TE = 31 ms, flip angle = 52°, field of view = 21 × 21 cm^2^).

Since our imaging protocol was similar to what was used in the human connectome project (HCP), our MRI and fMRI data were preprocessed by using the minimal preprocessing pipeline established for the HCP (using software packages AFNI, FMRIB Software Library, and FreeSurfer pipeline). After preprocessing, the images from individual subjects were co-registered onto a common cortical surface template (see details in^55^). Then the fMRI data were spatially smoothed by using a gaussian surface smoothing kernel with a 2mm standard deviation.

For each subject, the voxel-wise fMRI signal was standardized (i.e. zero mean and unitary standard deviation) within each session and was averaged across repeated sessions. Then the fMRI data were concatenated across different sessions and subjects for training the encoding model.

### Modeling and sampling the semantic space

To represent words as vectors, we used a pre-trained word2vec model^21^. Briefly, this model was a shallow neural network trained to predict the neighboring words of every word in the Google News dataset, including about 100 billion words (https://code.google.com/archive/p/word2vec/). After training, the model was able to convert any English word to a vector embedded in a 300-dimensional semantic space (extracted through software package Gensim^56^ in python). Note that the basis functions learned with word2vec should not be interpreted individually, but collectively as a space. Arbitrary rotation of the semantic space would end up with an equivalent space, even though it may be spanned by different semantic features. The model was also able to extract the semantic relationship between words by simple vector operations^30^. Individual words were extracted from audio stories using Speechmatics (https://www.speechmatics.com/), and then were converted to vectors through word2vec.

### Voxel-wise encoding model

We mapped the semantic space, as modeled by word2vec, to the cortex through voxel-wise linear encoding models, as explored in previous studies^1,24,38,39^. For each voxel, we modeled its response to a word as a linear combination of the word features in the semantic space.

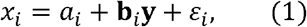

where *x_i_* is the response at the i-th voxel, **y** is the word embedding represented as a 300-dimensional column vector with each element corresponding to one axis (or feature) in the semantic space, **b**_*i*_ is a row vector of regression coefficients, *a_i_* is the bias term, and *ε_i_* is the error or noise.

### Training the encoding model

We used the (word, data) samples from the training stories to estimate the encoding model. As words occurred sequentially in the audio story, each word was given a duration based on when it started and ended in the audio story. A story was represented by a time series of word embedding sampled every 0.1 second. For each feature in the word embedding, its time-series signal was further convolved with a canonical hemodynamic response function (HRF) to account for the temporal delay and smoothing due to neurovascular coupling^57^. The HRF-convolved feature-wise representation was standardized and downsampled to match the sampling rate of fMRI.

It follows that the response of the i-th voxel at time t was expressed as Equation (2)

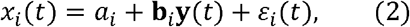

We estimated the coefficients (*a_i_*, **b**_*i*_) given time samples of (*x_i_*, **y**) by using least-squares estimation with L2-norm regularization. That is, to minimize the following loss function defined separately for each voxel.

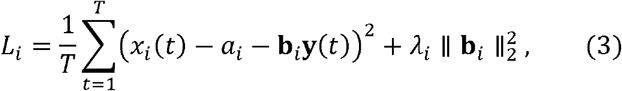

where *T* is the number of temporal samples, and λ_*i*_ is the regularization parameter for the i-th voxel.

We applied 10-fold generalized cross-validation^25^ in order to determine the regularization parameter. Specifically, the training data were divided evenly into 10 subsets, of which nine were used for model estimation and one was used for model validation. The validation was repeated 10 times such that each subset was used once for validation. In each time, the correlation between the predicted and measured fMRI responses was calculated and used to evaluate the validation accuracy. The average validation accuracy across all 10 times was considered as the cross-validation accuracy. We chose the optimal regularization parameter that yielded the highest cross-validation accuracy. Then we used the optimized regularization parameter and all training data for model estimation, ending up with the finalized model parameters denoted as 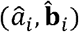.

### Cross validating the encoding model

We further tested the statistical significance of 10-fold cross-validation for every voxel based on a blockwise permutation test^58^. Specifically, we divided the training data into blocks; each block had a 20-sec duration. We kept the HRF-convolved word features intact within each block but randomly shuffled the block sequence for each of 100,000 trials of permutation. Before or after the block-wise shuffling, the word feature time series had the nearly identical magnitude spectrum, whereas the shuffling disrupted any word-response correspondence. For every trial of permutation, we ran the 10-fold cross-validation as aforementioned, resulting in a null distribution that included 100,000 cross-validation accuracies with permutated data. Against this null distribution, we compared the cross validation without permutation and calculated the one-sided p-value while testing the significance with FDR *q* < 0.05.

Following this 10-fold cross-validation, the model had been validated against 5,228 unique words. Thus, at the voxels of statistical significance, the word-evoked responses were considered to be predictable by the encoding models. Using the voxels of significance, we further created a cortical mask and confined the subsequent analyses to voxels in the created mask.

### Testing the encoding model

We also tested how well the encoding model could be generalized to a new story never used for model training and further evaluated how different regions varied in their responses to the same input stimuli. For this purpose, the trained encoding model was applied to the testing story, generating a voxel-wise model prediction of the fMRI response to the testing story.

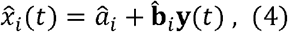

where **y**(*t*) is the HRF-convolved time series of word embedding extracted from the testing story.

To evaluate the encoding performance, we calculated the correlation between the predicted fMRI response 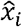 and the actually measured fMRI response *x_i_*. To evaluate the statistical significance, we used a block-wise permutation test^58^ (20-sec window size; 100,000 permutations) with FDR *q* < 0.05, similar to the analysis for cross-validation.

Since the measured fMRI responses to the testing story were averaged across sessions and subjects, the average responses had a much higher signal to noise ratio, or lower noise ceiling^59^, allowing us to visually inspect the encoding performance based on the response time series and to exam the response variation across regions. However, the testing story only included 368 unique words. Where the model succeeded in predicting the voxel response to the testing story was expected to be incomplete, relative to where the model would be able to predict given a larger set of word samples, e.g. as those used for cross validation.

In addition, we extracted the fMRI responses at ROIs predefined in the Human Brainnetome Atlas, which is a connectivity-based parcellation reported in an independent study^27^. We averaged the measured and model-predicted fMRI responses within each given ROI, and compared them as time series (see Figure 3c). The corresponding statistics regarding the location, size, and prediction performance of each ROI are listed in Supplementary Table 2.

### Mapping cortical representation of semantic categories

Using it as a predictive model, we further applied the estimated encoding model to a large vocabulary set including about 40,000 words^29^. At each voxel, we calculated the model-predicted response to every word and estimated the mean and the standard deviation for the response population and normalized the model-predicted response to any word as a z value.

Then we focused on the model prediction given 9,849 words from nine categories: *tool*, *human, plant, animal, place, communication, emotion, change, quantity* (Supplementary Table 3). See Supplementary Method 3 for more details about collecting samples for semantic categories. Every word had been rated for concreteness, ranging from 1 (most abstract) to 5 (most concrete). For each word, we used word2vec to compute its vector representation, and then used the voxel-wise encoding model to map its cortical representation.

As words were grouped by categories, we sought the common cortical representation shared by those in the same category. For this purpose, we averaged the cortical representation of every word in each category, and thresholded the average representation based on its statistical significance (one-sample t-test, FDR *p* < 0.01). We evaluated whether a given category was differentially represented by the left vs. right hemisphere, by counting for each hemisphere the number of voxels associated with that category. We also evaluated the semantic selectivity of each voxel, i.e. how the voxel was more selective to one category than the others. For a coarse measure of categorical selectivity, we identified, separately for each voxel of significance, a single category that resulted in the strongest voxel response among all nine categories and associated that voxel with the identified category (or by “winners take all”).

### Assessing word relations in the semantic space

Vector representations of words obtained by word2vec allow word relations to be readily extracted and applied with simple vector arithmetic^30^. For example, an arithmetic expression of (*hand* - *finger* + *second* leads to a vector close to that of *hour* in terms of cosine similarity. In this example, the subtraction extracts the relationship between *hand* and *finger*, which is intuitively interpretable as a *whole-part* relationship as a *finger* is part of a *hand*. It follows that the addition transfers this *whole-part* relationship to another word *second*, ending up with the word *hour*, while a *second* is indeed part of an *hour*.

Beyond this illustrative example, we examined a number of word pairs that held 1 out of 10 classes of semantic relations, as defined in or derived from the SemEval-2012 Task 2 dataset^31^. In this dataset, individual word-pair samples were scored by humans (by crowdsourcing) in terms of the degree to which a word pair could be viewed as an illustrative example of a specific semantic relation. The score ranged from −100 to 100 with −100 being the least and 100 being the most illustrative. For each class of semantic relation, we only included those word pairs with positive scores such that the included word pairs were affirmative samples that matched human understandings in a population level. We excluded the *reference* relation and separated the *space-time* relation into *space-associated* and *time-associated* relations. In brief, the 10 semantic relations (and their sample sizes) were *whole-part* (178 pairs), *class-inclusion* (113), *object-attribute* (63), *case relations* (106), *space-associated* (58), *time-associated* (44), *similar* (160), *contrast* (162), *object-nonattribute* (69), *cause-effect* (107). See details in Supplementary Table 4. Also see Supplementary Method 3 for details about collecting samples of semantic relations.

We investigated how generalizable semantic relations could be represented by differential vectors in the semantic space by using a leave-one-out test for each class of semantic relation. Specifically, we used the differential vector between any pair of words as the vector representation of their relation (or the “relation vector”). For a given class of semantic relation, we calculated the cosine similarity between the relation vector of every word-pair in the class and the average relation vector of all other word pairs in the same class (or the “matched relational similarity”) and compared it against the cosine similarity with the average relation vector in a different class (or the “unmatched relational similarity”). The matched relational similarity indicated the consistency of relation vectors in the same class of semantic relation. Its contrast against the unmatched relational similarity was evaluated with paired t-test (FDR *q* < 0.001). See more details in Supplementary Method 4 and the related results in Supplementary Figure 6.

### Mapping cortical representation of semantic relation

Applying the encoding model to the differential vector of a word pair could effectively generate the cortical representation of the corresponding word relation. With this notion, we used the encoding model to predict the cortical representations of semantic relations. For each class of semantic relation, we calculated the relation vector of every word pair in that class, projected the relation vector onto the cortex using the encoding model, and averaged the projected patterns across word-pair samples in the class. For the averaged cortical projection, we tested the statistical significance for every voxel based on a paired permutation test. In this test, we flipped every word pair at random for 100,000 trials. For every trial, we calculated the model-projected cortical pattern averaged across the randomly flipped word pairs, yielding a null distribution per voxel. Against this voxel-wised null distribution, we compared the average voxel value projected from non-flipped word pairs and calculated the two-sided p-value with the significance level at FDR *q* < 0.05. The resulting pattern of significant voxels was expected to report the primary cortical representation of each semantic relation of interest.

Complementary to the voxel-wise univariate analysis, we also applied a multivariate pattern analysis (MVPA) to the cortical projection of word relations^60^. Unlike the univariate analysis, MVPA was able to account for interactions between voxels and uncover likely multiple cortical patterns associated with each semantic relation of interest. Specifically, given a class of semantic relation, we concatenated the cortical pattern projected from every word-pair samples in that class and calculated a covariance matrix describing the similarity of representations between samples^61,62^. By using principal component analysis (PCA), we obtained a set of orthogonal components (i.e. eigenvectors), each representing a cortical pattern that accounted for the covariance to a decreasing extent. We chose the top-10 principal components and calculated the pattern-wise cosine similarity between every component and the cortical projection of every word-pair sample. For each component, we averaged the cosine similarity across all samples of the given semantic relation and tested the statistical significance based on one-sample t-test (*p* < 0.01). Specifically, for any relation with multiple significant components, we grouped and sorted the word-pairs based on their corresponding cosine similarities with each component. For each component, we listed the top-4 word-pairs with the highest cosine similarity in order to gain intuitive understanding as to whether the component was selective to a sub-class of that relation. See more details in Supplementary Method 7.

## Data availability

The raw and processed imaging datasets, as well as the supplementary data that support the findings of this study, are shared via a public repository in the Open Science Framework (https://osf.io/eq2ba/). The DOI of this dataset is 10.17605/OSF.IO/EQ2BA. The raw imaging datasets will also be shared via the OpenNeuro platform (https://openneuro.org/).

## Code availability

The code for training and testing the voxel-wise encoding model is also shared via the public repository in the Open Science Framework (https://osf.io/eq2ba/).

## Acknowledgment

The authors thank David Kemmerer for helpful discussions and comments to this paper. This work was supported by National Institute of Mental Health R01MH104402, Purdue University, and the University of Michigan.

## Author contributions

Z.L., Y.Z., R.W. designed research; Y.Z. and K.H. performed research; Y.Z. analyzed data; Y.Z. and Z.L. wrote the paper.

## References

1 Huth, A. G., de Heer, W. A., Griffiths, T. L., Theunissen, F. E. & Gallant, J. L. Natural speech reveals the semantic maps that tile human cerebral cortex. Nature 532, 453 (2016).

2 Yee, E. & Thompson-Schill, S. L. Putting concepts into context. Psychonomic bulletin & review 23, 1015–1027 (2016).

3 Holyoak, K. J. Analogy and relational reasoning. The Oxford handbook of thinking and reasoning, 234–259 (2012).

4 Mirman, D., Landrigan, J.-F. & Britt, A. E. Taxonomic and thematic semantic systems. Psychological bulletin 143, 499 (2017).

5 Bauer, A. J. & Just, M. A. Monitoring the growth of the neural representations of new animal concepts. Human brain mapping 36, 3213–3226 (2015).

6 Binder, J. R., Desai, R. H., Graves, W. W. & Conant, L. L. Where is the semantic system? A critical review and meta-analysis of 120 functional neuroimaging studies. Cerebral Cortex 19, 2767–2796 (2009).

7 Ralph, M. A. L., Jefferies, E., Patterson, K. & Rogers, T. T. The neural and computational bases of semantic cognition. Nature Reviews Neuroscience 18, 42 (2017).

8 Binder, J. R. In defense of abstract conceptual representations. Psychonomic bulletin & review 23, 1096–1108 (2016).

9 Patterson, K. & Ralph, M. A. L. in Neurobiology of Language 765–775 (Elsevier, 2015).

10 Humphreys, G. F., Hoffman, P., Visser, M., Binney, R. J. & Ralph, M. A. L. Establishing task-and modality-dependent dissociations between the semantic and default mode networks. Proceedings of the National Academy of Sciences, 201422760 (2015).

11 Martin, A. The representation of object concepts in the brain. Annu. Rev. Psychol. 58, 25–45 (2007).

12 Kiefer, M. & Pulvermüller, F. Conceptual representations in mind and brain: theoretical developments, current evidence and future directions, cortex 48, 805–825 (2012).

13 Mahon, B. Z. & Caramazza, A. Concepts and categories: A cognitive neuropsychological perspective. Annual review of psychology 60, 27–51 (2009).

14 Martin, A. GRAPES—Grounding representations in action, perception, and emotion systems: How object properties and categories are represented in the human brain. Psychonomic bulletin & review 23, 979–990 (2016).

15 Barsalou, L. W. On staying grounded and avoiding quixotic dead ends. Psychonomic bulletin & review 23, 1122–1142 (2016).

16 Sachs, O. et al. Automatic processing of semantic relations in fMRI: neural activation during semantic priming of taxonomic and thematic categories. Brain research 1218, 194–205 (2008).

17 Schwartz, M. F. et al. Neuroanatomical dissociation for taxonomic and thematic knowledge in the human brain. Proceedings of the National Academy of Sciences 108, 8520–8524 (2011).

18 Pulvermüller, F. How neurons make meaning: brain mechanisms for embodied and abstract-symbolic semantics. Trends in cognitive sciences 17, 458–470 (2013).

19 Hagoort, P. Nodes and networks in the neural architecture for language: Broca’s region and beyond. Current opinion in Neurobiology 28, 136–141 (2014).

20 Hasson, U., Malach, R. & Heeger, D. J. Reliability of cortical activity during natural stimulation. Trends in cognitive sciences 14, 40–48 (2010).

21 Mikolov, T., Sutskever, I., Chen, K., Corrado, G. S. & Dean, J. in Advances in neural information processing systems. 3111–3119.

22 Mitchell, T. M. et *al*. Predicting human brain activity associated with the meanings of nouns. science 320, 1191–1195 (2008).

23 Anderson, A. J. et al. Predicting neural activity patterns associated with sentences using a neurobiologically motivated model of semantic representation. Cerebral Cortex 27, 4379–4395 (2016).

24 Naselaris, T., Kay, K. N., Nishimoto, S. & Gallant, J. L. Encoding and decoding in fMRI. Neuroimage 56, 400–410 (2011).

25 Golub, G. H., Heath, M. & Wahba, G. Generalized cross-validation as a method for choosing a good ridge parameter. Technometrics 21, 215–223 (1979).

26 Knecht, S. et al. Language lateralization in healthy right-handers. Brain 123, 74–81 (2000).

27 Fan, L. et al. The human brainnetome atlas: a new brain atlas based on connectional architecture. Cerebral cortex 26, 3508–3526 (2016).

28 Miller, G. WordNet: An electronic lexical database. (MIT press, 1998).

29 Brysbaert, M., Warriner, A. B. & Kuperman, V. Concreteness ratings for 40 thousand generally known English word lemmas. Behavior research methods 46, 904–911 (2014).

30 Mikolov, T., Yih, W.-t. & Zweig, G. in Proceedings of the 2013 Conference of the North American Chapter of the Association for Computational Linguistics: Human Language Technologies. 746–751.

31 Jurgens, D. A., Turney, P. D., Mohammad, S. M. & Holyoak, K. J. in Proceedings of the First Joint Conference on Lexical and Computational Semantics-Volume 1: Proceedings of the main conference and the shared task, and Volume 2: Proceedings of the Sixth International Workshop on Semantic Evaluation. 356–364 (Association for Computational Linguistics).

32 Raichle, M. E. et al. A default mode of brain function. Proceedings of the National Academy of Sciences 98, 676–682 (2001).

33 Corbetta, M. & Shulman, G. L. Control of goal-directed and stimulus-driven attention in the brain. Nature reviews neuroscience 3, 201 (2002).

34 Scolari, M., Seidl-Rathkopf, K. N. & Kastner, S. Functions of the human frontoparietal attention network: Evidence from neuroimaging. Current opinion in behavioral sciences 1, 32–39 (2015).

35 Pennington, J., Socher, R. & Manning, C. in Proceedings of the 2014 conference on empirical methods in natural language processing (EMNLP). 1532–1543.

36 Pereira, F., Gershman, S., Ritter, S. & Botvinick, M. A comparative evaluation of off-the-shelf distributed semantic representations for modelling behavioural data. Cognitive neuropsychology 33, 175–190 (2016).

37 Huth, A. G., Nishimoto, S., Vu, A. T. & Gallant, J. L. A continuous semantic space describes the representation of thousands of object and action categories across the human brain. Neuron 76, 1210–1224 (2012).

38 Wen, H. et al. Neural encoding and decoding with deep learning for dynamic natural vision. Cerebral Cortex, 1–25 (2017).

39 Pereira, F. et al. Toward a universal decoder of linguistic meaning from brain activation. Nature communications 9, 963 (2018).

40 Rubenstein, H. & Goodenough, J. B. Contextual correlates of synonymy. Communications of the ACM3, 627–633 (1965).

41 Vincent-Lamarre, P. et al. The latent structure of dictionaries. Topics in cognitive science 8, 625–659 (2016).

42 Binder, J. R. et al. Toward a brain-based componential semantic representation. Cognitive neuropsychology 33, 130–174 (2016).

43 Binder, J. R., Westbury, C. F., McKiernan, K. A., Possing, E. T. & Medler, D. A. Distinct brain systems for processing concrete and abstract concepts. Journal of cognitive neuroscience 17, 905–917 (2005).

44 Wang, J., Conder, J. A., Blitzer, D. N. & Shinkareva, S. V. Neural representation of abstract and concrete concepts: A meta-analysis of neuroimaging studies. Human brain mapping 31, 1459–1468 (2010).

45 Bunge, S. A., Wendelken, C., Badre, D. & Wagner, A. D. Analogical reasoning and prefrontal cortex: evidence for separable retrieval and integration mechanisms. Cerebral cortex 15, 2392–49 (2004).

46 Smith, S. M. et al. Correspondence of the brain’s functional architecture during activation and rest. Proceedings of the National Academy of Sciences 106, 13040–13045 (2009).

47 Fox, M. D. et al. The human brain is intrinsically organized into dynamic, anticorrelated functional networks. Proceedings of the National Academy of Sciences of the United States of America 102, 9673–9678 (2005).

48 Andrews-Hanna, J. R., Reidler, J. S., Sepulcre, J., Poulin, R. & Buckner, R. L. Functional-anatomic fractionation of the brain’s default network. Neuron 65, 550–562 (2010).

49 Spreng, R. N. The fallacy of a “task-negative” network. Frontiers in psychology 3, 145 (2012).

50 Simony, E. et al. Dynamic reconfiguration of the default mode network during narrative comprehension. Nature communications 7, 12141 (2016).

51 Sormaz, M. et al. Default mode network can support the level of detail in experience during active task states. Proceedings of the National Academy of Sciences 115, 9318–9323 (2018).

52 Spunt, R. P., Kemmerer, D. & Adolphs, R. The neural basis of conceptualizing the same action at different levels of abstraction. Social cognitive and affective neuroscience 11, 1141–1151 (2015).

53 Gotts, S. J. et al. Two distinct forms of functional lateralization in the human brain. Proceedings of the National Academy of Sciences 110, E3435–E3444 (2013).

54 Maaten, L. v. d. & Hinton, G. Visualizing data using t-SNE. Journal of machine learning research 9, 2579–2605 (2008).

55 Glasser, M. F. et al. The minimal preprocessing pipelines for the Human Connectome Project. Neuroimage 80, 105–124 (2013).

56 Rehurek, R. & Sojka, P. in In Proceedings of the LREC 2010 Workshop on New Challenges for NLP Frameworks. (Citeseer).

57 Lindquist, M. A., Loh, J. M., Atlas, L. Y. & Wager, T. D. Modeling the hemodynamic response function in fMRI: efficiency, bias and mis-modeling. Neuroimage 45, S187–S198 (2009).

58 Adolf, D. et al. Increasing the reliability of data analysis of functional magnetic resonance imaging by applying a new blockwise permutation method. Frontiers in neuroinformatics 8, 72 (2014).

59 Sahani, M. & Linden, J. F. in Advances in neural information processing systems. 125–132.

60 Haxby, J. V. Multivariate pattern analysis of fMRI: the early beginnings. Neuroimage 62, 852–855 (2012).

61 Kriegeskorte, N., Mur, M. & Bandettini, P. A. Representational similarity analysis-connecting the branches of systems neuroscience. Frontiers in systems neuroscience 2, 4 (2008).

62 Lazar, N. The statistical analysis of functional MRI data. (Springer Science & Business Media, 2008).

