## Supplementary Information for "Connecting Concepts in the Brain by Mapping Cortical Representations of Semantic Relations"

Zhang et al.

### Supplementary Methods

#### 1. Counterbalancing stories across subjects

In order to thoroughly sample the semantic space, we designed experiments to have each participant listen to different stories during fMRI and used the obtained stimulus-response samples to train and cross-validate the encoding model. By counterbalancing the stories across subjects, we aimed to sample different words with different subjects, while having the sampled words cover similar distributions in the semantic space, include a similar set of frequent words, and sample every semantic category or relation of interest in nearly identical proportions across individual subjects.

In the following, we refer to the stories presented to each subject as the "subject-specific story set". Each subject-specific story set contained 2 or 3 stories and 2,492 ± 423 words. When combined across subjects, the story sets sampled a total of 47,356 words, including 5,228 unique words. We considered this sample size sufficiently large because it outnumbered the English words for daily use (around 3,000).

Among the words included for model training, 50 words were most frequent, including "*I*" (2,464 times), "*the*" (2,074), "*was*" (952), "*my*" (912), "*that*" (889), "*in*" (781), "*you*" (753), "*it*" (663), "*so*" (598), "*me*" (531), "*he*" (480), "*like*" (472), "we" (433), "*she*" (428), "*on*" (425). For each of these frequent words, its occurrences were about evenly distributed across subjects (Supplementary Figure 2). Since each story we chose was a natural story told by a storyteller in a live talk (Moth Radio Hour), the frequency of words we sampled in the entire experiment was generally close to the frequency of words as used in daily life. See Supplementary Table 1 for more details about the top-5 frequent words in the stories presented to each subject.

We visualized the distribution of subject-specific word samples in the semantic space. For the sake of illustration, the semantic space was reduced to a 2-D plane by applying nonlinear dimension reduction based on t-distributed stochastic neighbor embedding (t-SNE) [1], which is a widely-used nonlinear dimensionality reduction algorithm. For each story in the training dataset, we first used Speechmatics[1] to extract individual words in the transcript and then converted words to vectors through word2vec. In this way, we converted 47,356 words in the whole training dataset to a set of 300-dimension vectors. We implemented t-SNE in Matlab, specifying the algorithm to maintain the pairwise cosine distance between words. This algorithm iteratively updated the two-dimensional points to minimize the Kullback-Leibler divergence between distributions of points in the original high dimensional space and the dimension-

[1] https://www.speechmatics.com/

reduced space. After this process, we obtained 47,356 points in a two-dimensional space, which could be visualized in a scatter plot. We further labeled words by subject to visually compare their distributions.

As shown in Supplementary Figure 1, the word samples for every subject had nearly uniform distributions in the semantic space and the distributions were nearly identical across subjects. Since we concatenated the whole training dataset for this analysis, all plots in Supplementary Figure 1 shared the same x-y axis. This result suggests that the words sampled with every subject were highly comparable in terms of their distribution in the semantic space without any notable sampling bias.

We also tested how individual semantic categories or relations were sampled across subjects in order to ensure that none of the subject-specific story sets was biased towards any semantic category or relation over others. For a given category, we counted the number of times that words in that category occurred in the stories presented to each subject and compared the word counts across subjects (Supplementary Figure 3). Similarly, for a given semantic relation, we counted the number of times that word pairs in that relation occurred in the stories presented to each subject and compared the resulting counts across subjects (Supplementary Figure 4). We found that each semantic category or relation of interest was about evenly sampled across subjects without any notable sampling bias. The fact that the stories used in this study were all naturalistic precluded us from ensuring precisely even and balanced sampling across subjects. Nevertheless, the subject-wise sampling bias was minor and unlikely to invalidate our method or significantly confound our findings.

#### 2. Testing effects of individual variance

Individual variation is unavoidable. For example, individuals vary in terms of laterality as demonstrated before [2,3]. In this study, we attempted to identify where individual variation was most pronounced in the brain with respect to cortical activation with naturalistic audio stories. For the purpose, we used the fMRI data when every subject was listening to the same testing story in two repeated sessions. For each session, we split the data into 4 non-overlapping segments (or sub-stories). For each segment, we first computed the voxel-wise correlation between the fMRI signals from the 2 repeated sessions in every subject. As a result, the z-transformed correlation at every voxel depended on two factors: subject & stimulus. We performed a two-way analysis of variance (ANOVA) and identified the voxels where the factor of the subject had a significant effect. In this way, we identified the cortical locations where activation was significantly different across subjects during natural speech comprehension.

As shown in Supplementary Figure 11, the effect of individual variation (with uncorrected $p < 0.01$) was found primarily in the auditory association cortex; however, the effect was largely diminished after correction for multiple comparisons (FDR $q < 0.05$). These results suggest that there was a minor degree of individual variation in cortical activation induced by audio stories, consistent with the findings from prior studies [4].

##### 3. Collecting samples of semantic categories and relations

In this study, we collected the word samples of different semantic categories from a large corpus (near 40 thousand words) including the most commonly used English words as documented in a previous paper [5].

For a specific semantic category, we identified a subset that fitted the synset (a cognitive synonym) of this category based on Wordnet[2] [6]. For example, the "tool" category was associated with the synset "*tool.n.01*" in Wordnet. This synset is defined as "an implement used in the practice of a vocation". For each semantic category, we listed its name, the name and definition of its corresponding Wordnet synset, the number of word samples, and example words in Supplementary Table 3.

We collected the word-pair samples of different semantic relationships from the SemEval-2012 task 2 dataset [7], which is downloadable from an online resource[3]. This dataset included 10 general relations (1. class-inclusion, 2. part-whole, 3. similar, 4. contrast, 5. attribute, 6. non-attribute, 7. case relations, 8. cause-purpose, 9. space-time, 10. reference). Each of these relations contains several subcategories of "finely defined relations". The detailed definitions of the 79 finely defined relations could be found in the document "subcategories-definitions.txt" [3]. Starting with those definitions, we redefined the 10 semantic relations for this study. Specifically, we excluded the "reference" relationship because its sub-classes were less consistent ("*sign:significant*", "*representation*", and "*knowledge*" etc.). We also separated the "space-time" relationship to "space-associated" and "time-associated" relations since these were intuitively two different semantic relations. We collected word-pair samples from "SemEval-2012-Platinum-Ratings" package – the latest release of the dataset. The dataset provided a human-rated score obtained from crowdsourcing and the score indicated the relevance between a word pair and a semantic relation. The score ranged from -100 to 100 with -100 being the least and 100 being the most illustrative. For each class of semantic relation, we only included those word pairs with positive scores such that the included word pairs were affirmative samples that matched human understandings about word/semantic relations. In the following, we listed the sets of finely defined relations from which we collected the

---

[2] https://wordnet.princeton.edu/
[3] https://sites.google.com/site/semeval2012task2/

word-pair samples for each semantic relation. The labels and definitions listed above are according to the original documents in the SemEval-2012 task 2 dataset.

The “**whole-part**” relationship was collected from the following finely identified relations: 2a (*object:component*), 2b (*collection:member*), 2c (*mass:potion*), 2d (*event:feature*), 2e (*activity:stage*), 2f (*item:topological part*), 2g (*object:stuff*), and 2h (*creature:possession*).

The “**class-inclusion**” relationship was collected from the following finely identified relations: 1a (*taxonomic*), 1b (*functional*), 1c (*singular collective*), 1d (*plural collective*), and 1e (*class:individual*).

The “**object-attribute**” relationship was collected from the following finely identified relations: 5a (*item:attribute*), 5c (*object:state*), and 5e (*object:typical action*).

The “**case relations**” was collected from the following finely identified relations: 7a (*agent:object*), 7b (*case relations*), 7c (*agent:instrument*), 7d (*action:object*), and 7e (*action:recipient*).

The “**space-associated**” relationship was collected from the following finely identified relations: 9b (*location:process/product*), 9c (*location:action/activity*), and 9d (*location:instrument/associated item*).

The “**time-associated**” relationship was collected from the following finely identified relations: 9f (*time action/activity*) and 9g (*time associated item*).

The “**similar**” relationship was collected from the following finely identified relations: 3a (*synonymity*), 3b (*dimensional similarity*), 3c (*dimensional excessive*), 3d (*dimensional naughty*), 3e (*conversion*), 3f (*attribute similarity*), 3g (*coordinates*), and 3h (*change*).

The “**contrast**” relationship was collected from the following finely identified relations: 4a (*contradictory*), 4b (*contrary*), 4c (*reverse*), 4d (*directional*), 4e (*incompatible*), 4f (*asymmetric contrary*), 4g (*pseudoantonym*), and 4h (*defective*).

The “**object-nonattribute**” relationship was collected from the following finely identified relations: 6a (*item:nonattribute*), 6c (*object:nonstate*), and 6e (*objects:atypical action*).

The “**cause-effect**” relationship was collected from the following finely identified relations: 8a (*cause:effect*), 8b (*cause:compensatory action*), 8d (*action/activity:goal*), 8e (*agent:goal*), and 8f (*instrument:goal*).

#### 4. Evaluating the consistency of relation vector within and between semantic relations

We would like to address whether the vector representation of a semantic relation is generalizable in the word2vec space. Results from a prior study [8] suggest that word2vec preserves the semantic and syntactic relations between words through vector arithmetic. We did a similar analysis with our dataset and showed consistent results to reinforce this conclusion. In the following, we elaborate our analysis.

For a pair of words, we represented their relationship as a vector, referred to as the relation vector, and calculated the relation vector by subtracting the vector representation of the second word from that of the first word. For two relation vectors, we evaluated their similarity, or relational similarity, based on their cosine similarity, as in the prior study [8]. For a semantic relation, we calculated and averaged the relation vector for every word-pair sample of that relation, and used the averaged relation vector as the vector representation of the semantic relation.

Given the above notation and calculation, we hypothesized that the relation vector of a given word pair was more (cosine) similar to the relation vector of the semantic relation that matched the relationship of that word pair than the relation vector of a semantic relation that did not match it. To test this hypothesis, we did a "leave-one-out" test described below.

We began by picking one semantic relation of interest, e.g. "whole-part", and then we chose one sample of that relation, e.g. <book, pages>, as the testing example. We calculated the relation vector of the testing example. Then we left this testing example out of the sample set for the corresponding relation, calculated the relation vectors of all remaining samples, and averaged them to obtain the relation vector of the semantic relation while excluding the contribution from the testing example. We calculated the similarity in the relation vector between the testing example and the semantic relation that matched that example, referred to as "matched relational similarity". The "matched relational similarity" indicated the degree to which the relation vector of a word pair was consistent with the relation vectors of other word pairs in the same class of semantic relation. For comparison, we also calculated the similarity in the relation vector between the testing example and a different semantic relation (one of the other 9 relations in this study), which did not match the testing example, referred to as "unmatched relational similarity". By choosing every word-pair sample once as the testing example, we obtained a set of samples for the "matched relational similarity", as well as a paired set of samples for the "unmatched relational similarity". We tested their difference and evaluated its statistical significance based on pair t-test (FDR $q < 0.001$). See results in Supplementary Figure 6.

In addition, we also performed a classification task. In this task, we classified each word pair (or the testing example) as one (out of 10) semantic relation, based on the relational similarity between the testing example and every semantic relation. Note that before classification, we excluded the testing example from the semantic relation it belonged to. The semantic relation that yielded the highest similarity was identified. The top-1 classification accuracy was calculated as a percentage and compared against the percentage by which the testing example was misclassified into a different semantic relation. See results in Supplementary Figure 7.

**5. Evaluating the association between semantic relations and semantic categories**

We noted that some semantic relations were naturally associated with specific semantic categories. For example, the "space-associated" relation tended to be associated with the "place" category. Such relation-category associations did not result from any intended bias by our data or design; instead they reflected natural language statistics for English.

Nevertheless, we evaluated the association between each semantic relation of interest and each semantic category of interest. Specifically, we counted how many words as included in the samples of a relation were also included in the samples of a category and divided this word count by the total number of word samples included in the relation. We did this separately for the first word and the last word and calculated the absolute difference as a measure of the association between a specific relation and a specific category, ranging from 0% to 100%.

See results in Supplementary Figure 10.

**6. Testing the effect of samples size**

As described in Supplementary Method 3, we maximized the sample size given the use of the materials used in this study [5]. However, the sample sizes were unequal for different categories. This unequal sampling did not result from any intended bias by our design but reflected the uneven categorization of English words. Nevertheless, we evaluated the effect of sample size on maps of cortical representations.

For each category, we randomly selected a subset of word samples (10% to 90%) and evaluated the corresponding cortical representations. For example, the maximum number of words in the "animal" category was 734, of which a subset was selected with the sample size ranging from 10% (73) to 100% (734) of the original sample size. We identified the voxels at which the model-predicted responses were significant for a varying number of words included in that category. Given a subset of word samples in one category, we averaged the cortical representation of every word in that set, and thresholded the average representation based on its statistical significance (one sample t-test, $FDR < 0.01$). We counted the total number of significant voxels in this average representation. We repeated this process for 100 independent trials given a proportion of word samples (10% to 90%) for each category, respectively. We then evaluated how the statistics (mean and standard error) of the total number of significant voxels changed as the sample size increases (Supplementary Figure 5).

#### 7. Mapping cortical representation of semantic relations using multivariate analysis

The univariate voxel-wise analysis was intended to address the cortical representation of semantic relations separately for individual voxels. However, a semantic relation could be represented by cortical networks that capture interaction across voxels, instead of by tuning properties of individual voxels. In line with this notion, we also conducted a multivariate analysis [9] by viewing the cortical projections of word relationships as pattern-wise responses, as opposed to voxel-wise responses.

Specifically, we applied a principal component analysis (PCA) to the covariance matrix that described the similarity of representations between word-pair samples. For each semantic relation of interest, we converted every word pair $k$ into two vectors and calculated their difference by element-wise subtraction. We then projected this vector difference in the semantic space to the brain through the encoding model. By doing so, we obtained a cortical pattern, denoted as $\mathbf{C}_k$, for the $k$-th word pair**.** We concatenated these cortical patterns to construct an $N$-by-$P$ matrix (where N is the number of word-pair samples and P is the number of voxels). To this matrix, we applied PCA and kept the first 10 principal components. These components in the brain space were multivariate spatial patterns (**SP**) that could be viewed as a set of orthogonal basis functions $\mathbf{SP}_1, \mathbf{SP}_2$ ...which explained the voxel interactions as cortical networks.

For each spatial pattern $\mathbf{SP}_i$**,** we then calculated its cosine similarity (denoted as $d_k^i$) with the cortical projection of every word-pair sample $k$.

$$d_k^i = < \frac{\mathbf{C}_k}{\|\mathbf{C}_k\|}, \mathbf{SP}_i >$$

Note that $\mathbf{SP}_i$ was the unitary vector and $d_k^i$ ranged from -1 to 1. For each pattern $\mathbf{SP}_i$, we obtained $N$ samples of $d_k^i$. By applying a one-sample t-test to these $N$ samples, we tested the significance of each spatial pattern ($p < 0.01$). If the cosine similarity with a spatial pattern had an average less than zero, we reversed the spatial pattern (multiplication by -1) such that the pattern was positively associated with the semantic relation. See results in Supplementary Figure 9. For a spatial pattern of statistical significance, we ranked the word pairs by $d_k^i$ and picked the top-4 most matched word pairs, in an attempt to probe its differential selectivity to various word relationships.

### Supplementary Figures

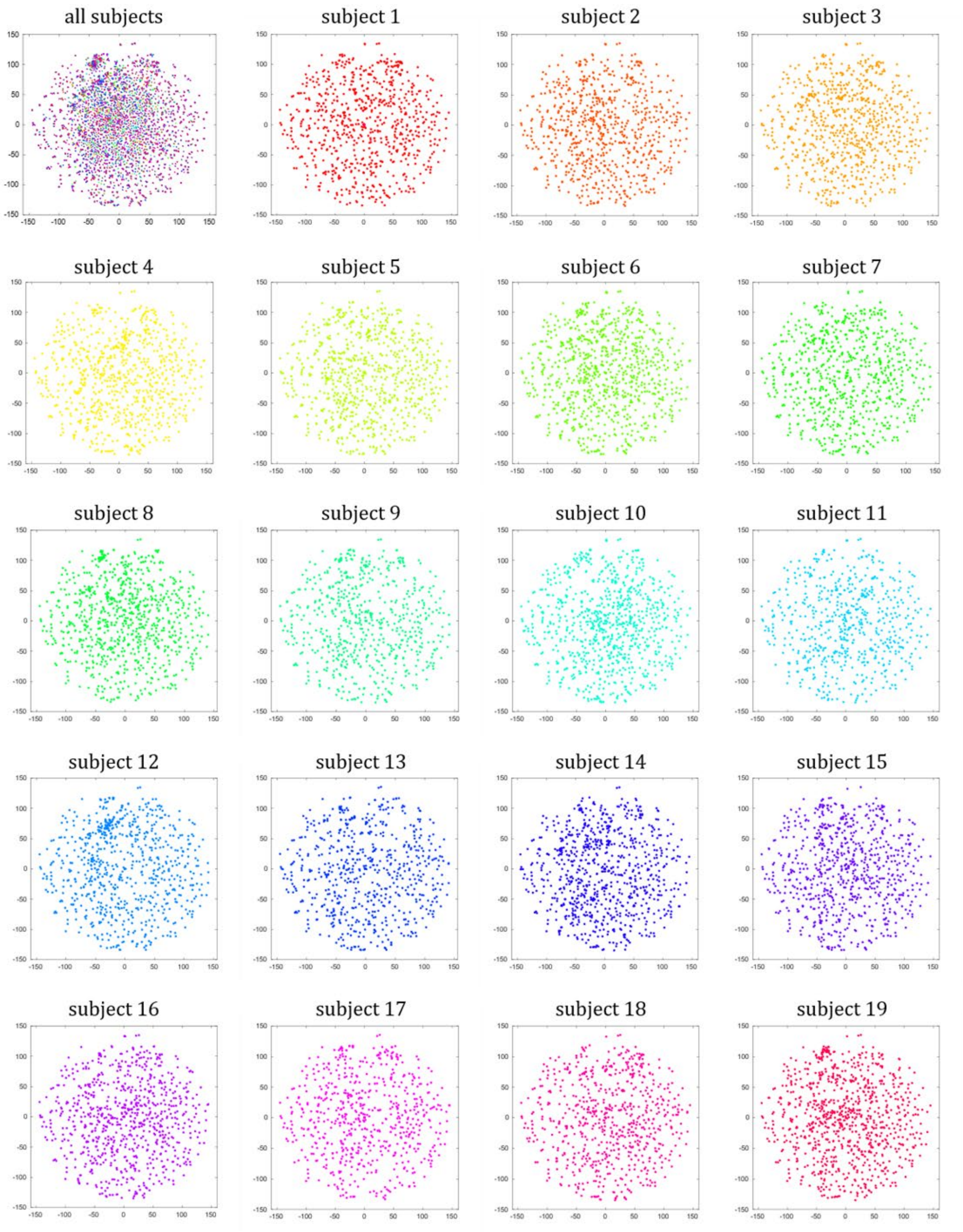


**Supplementary Figure 1. Visualization of word distributions in the training dataset through t-SNE plots.** For every word in the training stories, we convert it to a word embedding through the pre-trained word2vec model [10] and represent it as a vector in the 300-dimensional semantic space. We use t-distributed Stochastic Neighbor Embedding (t-SNE) [1] algorithm in Matlab to visualize the distribution of all words in the training dataset by reducing the embedding dimension while keeping the relevant pair-wise cosine similarity. We label every word by subject, for which the word is used as a stimulus. Each subject is coded by a different color. All panels share the same x-y axis, which is modeled by t-SNE on the training data concatenated from all subjects. Separate panels for individual subjects show that word sample distributions are roughly uniformly distributed in the semantic space. See Supplementary Method 1 for more details about counterbalancing stories across subjects.

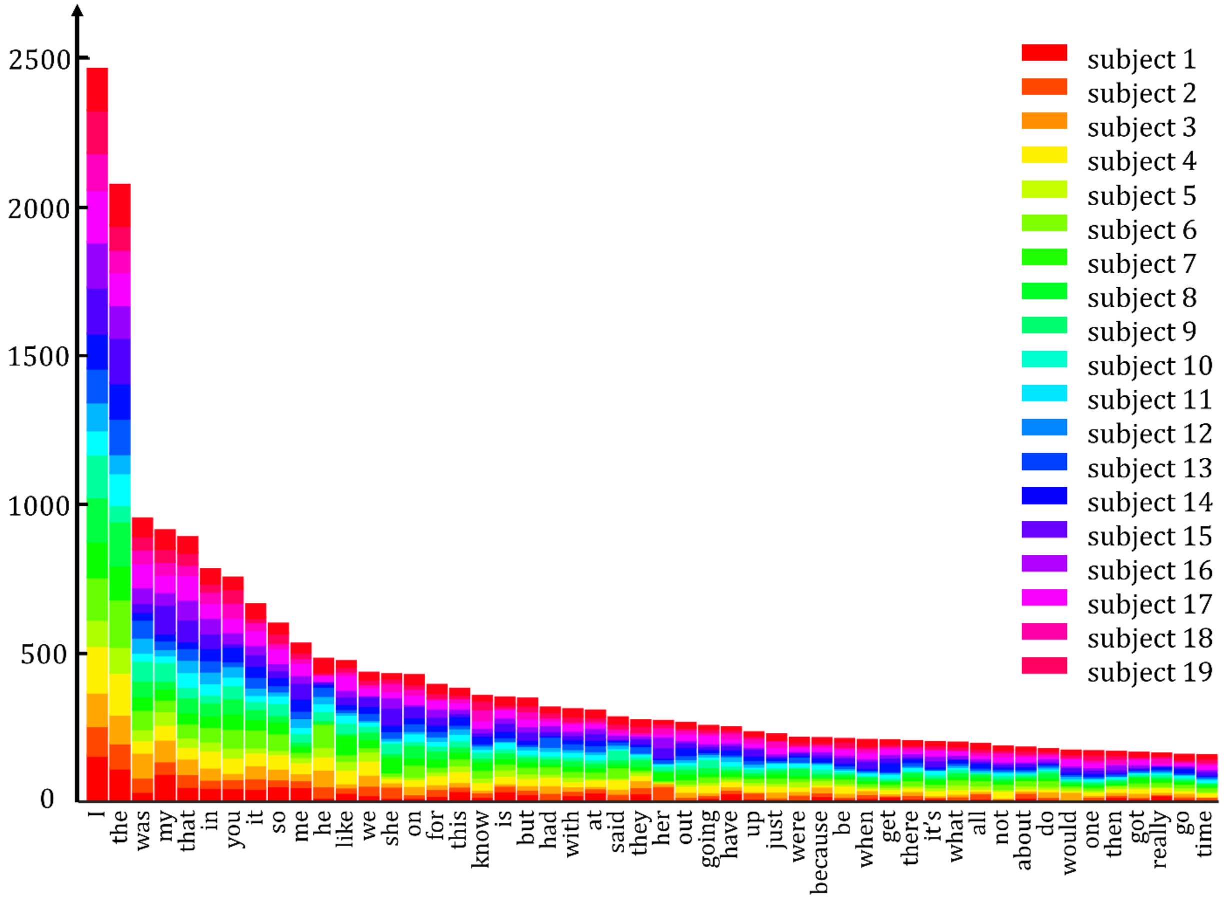


**Supplementary Figure 2. Distribution of top 50 frequent words in the training stories.** We count the frequency of each word in the training stories and sort every word in descending order of frequency (See Supplementary Method 1). The size of word samples is 47,356. Words that most frequently appear in the training stories include "*I*" (2,464 times), "*the*" (2,074), "*was*" (952), "*my*" (912), "*that*" (889), "*in*" (781), "*you*" (753), "*it*" (663), "*so*" (598), "*me*" (531), "*he*" (480), "*like*" (472), "*we*" (433), "*she*" (428), "*on*" (425). For each of these frequent words, we count its occurrences across subjects (as color-coded), as indicated by the height of each color-coded section. See Supplementary Method 1 for more details about counterbalancing stories across subjects.

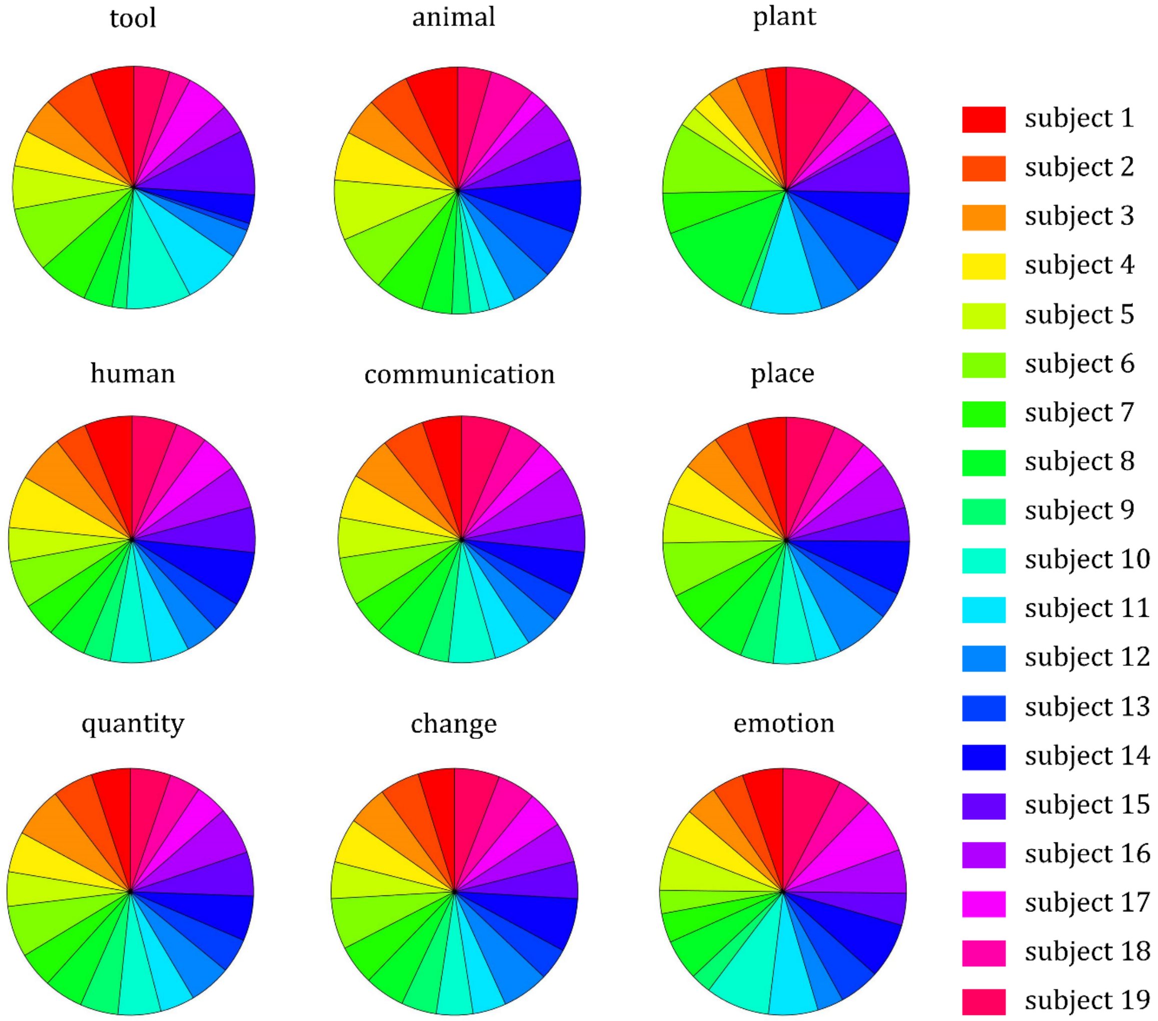


| Category | tool | animal | plant | human | communi cation | place | quantity | change | emotion |
|---|---|---|---|---|---|---|---|---|---|
| ① | 173 / 32 | 642 / 123 | 122 / 50 | 398 / 80 | 2159 / 251 | 5975 / 484 | 487 / 96 | 7346 / 703 | 7431 / 342 |
| ② | 200 | 734 | 387 | 808 | 2027 | 814 | 958 | 3417 | 504 |

**Supplementary Figure 3. Distribution of category-specific words across subjects.** We count the number of times that words in each category occurred in the training stories used for each subject (color-coded). The pie chart displays the proportion that category-specific words are distributed across subjects. In the table, row ① shows the total number of (repeated or unique) word occurrences summed across subjects; row ② shows the total number of word samples to which the encoding model is applied for mapping each category. See Supplementary Method 1 for more details about counterbalancing stories across subjects.

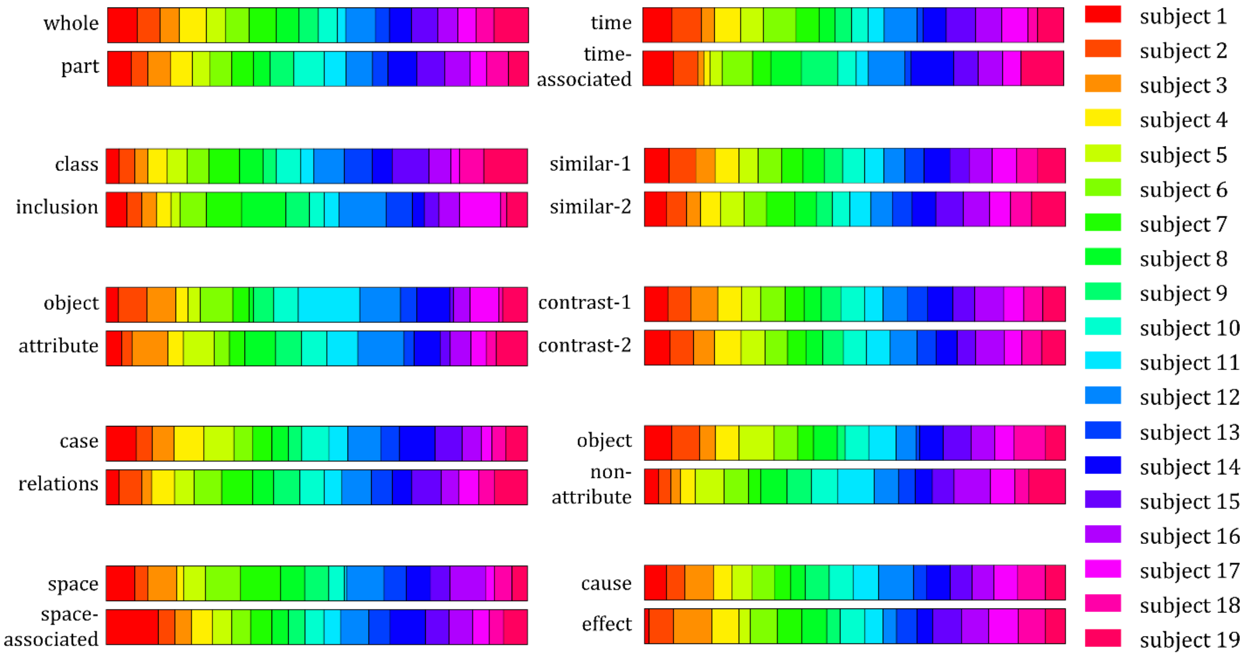


**Supplementary Figure 4. Distribution of relation-specific words across subjects.** We count the number of times that words in each relation occur in the training stories used for each subject (color-coded). We count this number separately for the first word (e.g. the "whole" word in the "whole-part" relation) and the second word (e.g. the "part" word in the "whole-part" relation). The bar chart displays the distribution of relation-specific word counts across subjects. The width of a color section indicates the proportion of words sampled for each subject. See Supplementary Method 1 for more details about counterbalancing stories across subjects.

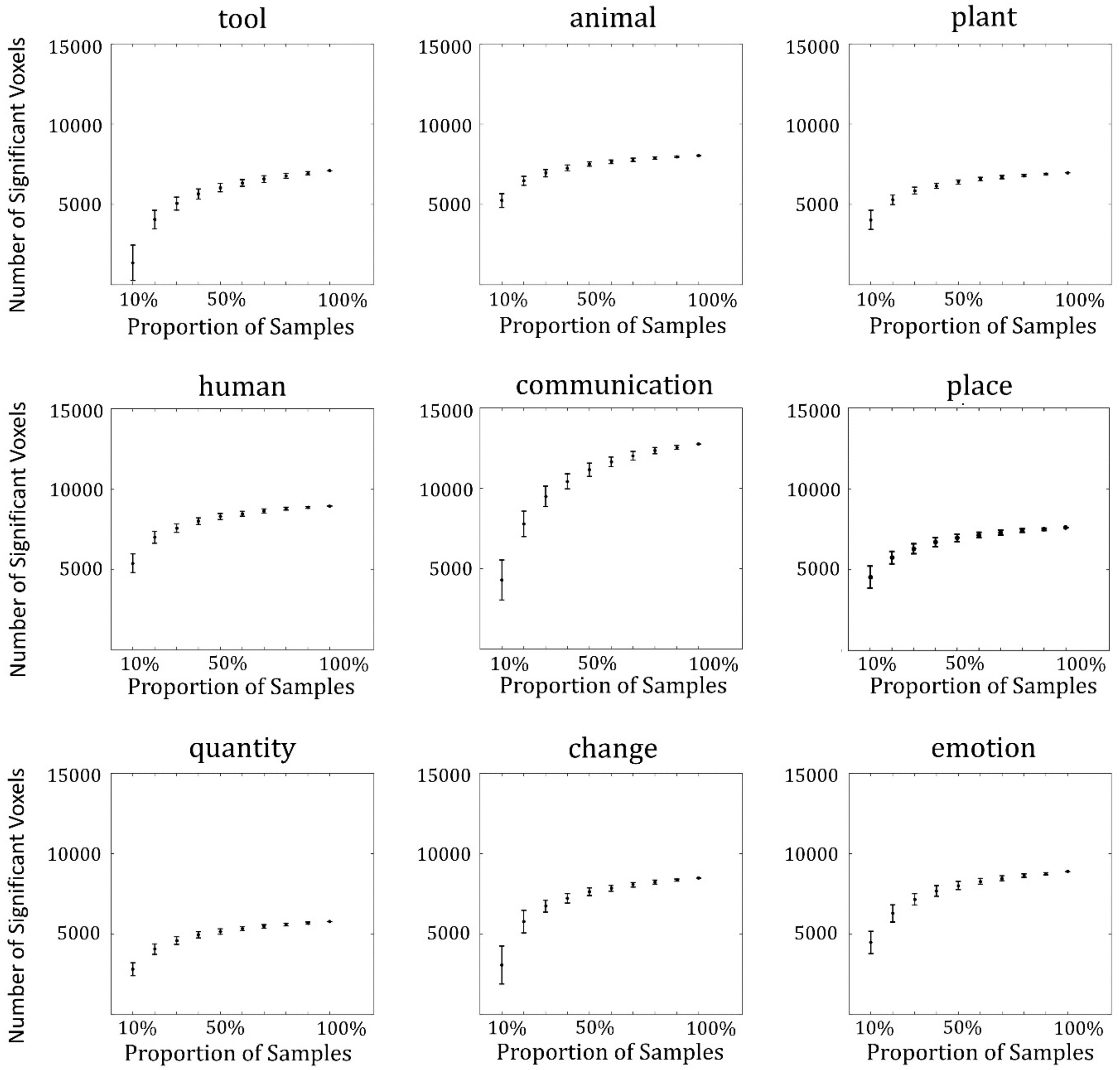


**Supplementary Figure 5. Effect of the sample size on cortical representation of each semantic category.** Each plot shows the number of significant voxels (one sample t-test, one-sided, FDR $q < 0.01$) as a function of the relative (%) sample size for each category. To get a robust estimation on the effect of sample size, we randomly sample a subset containing 10%, 20%, …, 90% of the samples, respectively, for 100 independent trials. The error bar indicates the standard error of the results across the 100 trials. See Supplementary Method 6 for more details about testing the effect of the sample size.

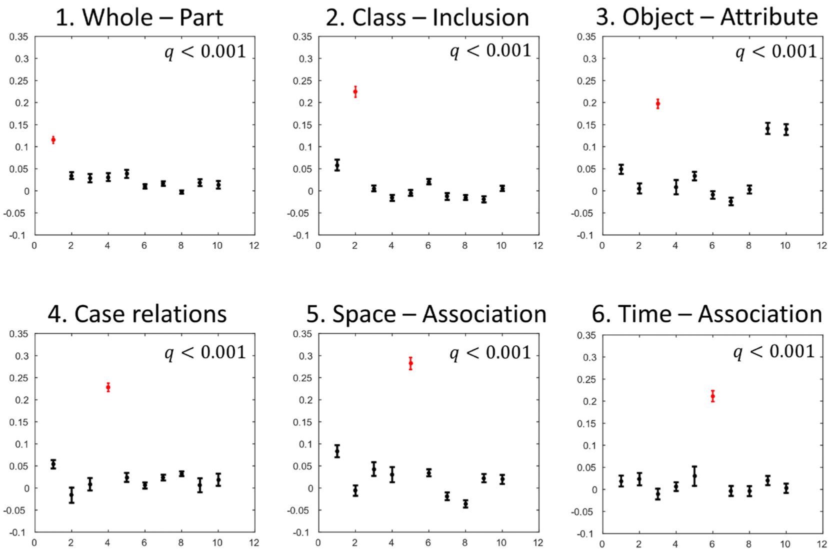


| Relation | M | U |
|---|---|---|
| 1 | 0.12 | 0.02 |
| 2 | 0.22 | 0.00 |
| 3 | 0.20 | 0.04 |
| 4 | 0.23 | 0.02 |
| 5 | 0.29 | 0.02 |
| 6 | 0.21 | 0.01 |
| 7 | 0.05 | 0.00 |
| 8 | 0.03 | 0.00 |
| 9 | 0.09 | 0.02 |
| 10 | 0.08 | 0.02 |

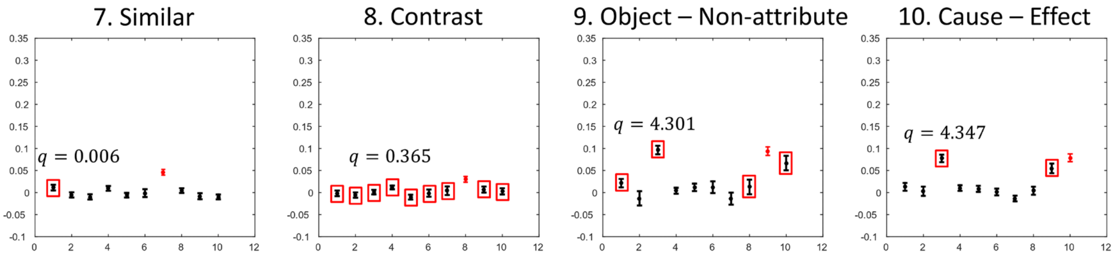


**Supplementary Figure 6. The matched vs. unmatched relational similarity in the semantic space.** Each plot shows the average "*matched*" relational similarity (red) or "*unmatched*" relational similarity (black). The relational similarity is calculated as the cosine similarity (between -1 and 1) between two vectors. The error bar indicates the standard error of the mean relational similarity across all word-pairs in a semantic relation. The red box indicates a lack of statistical significance between the "matched" vs. "unmatched" relational similarity (paired t-test, one-sided, FDR $q < 0.001$). The q-values in the box indicated the largest q-value among the "unmatched" relational similarity. For relations *whole-part*, *class-inclusion*, *object-attribute*, *case-relations*, *space-association*, and *time-association*, all "unmatched" relational similarity is significantly smaller than "matched" similarity with $q < 0.001$. However, for relations *similar* ("unmatched" *whole-part*, $q = 0.006$), *contrast* ("unmatched" *case relations*, $q = 0.365$), *object-nonattribute* ("unmatched" *object-attribute*, $q = 4.301$), and *cause-effect* ("unmatched" *object-attribute*, $q = 4.347$), the "matched" relational similarity is not significantly larger than all "unmatched" similarity. The table displays the "*matched*" relational similarity (column M) and the "*unmatched*" relational similarity on average (column U). See Supplementary Method 4 for more analysis details.

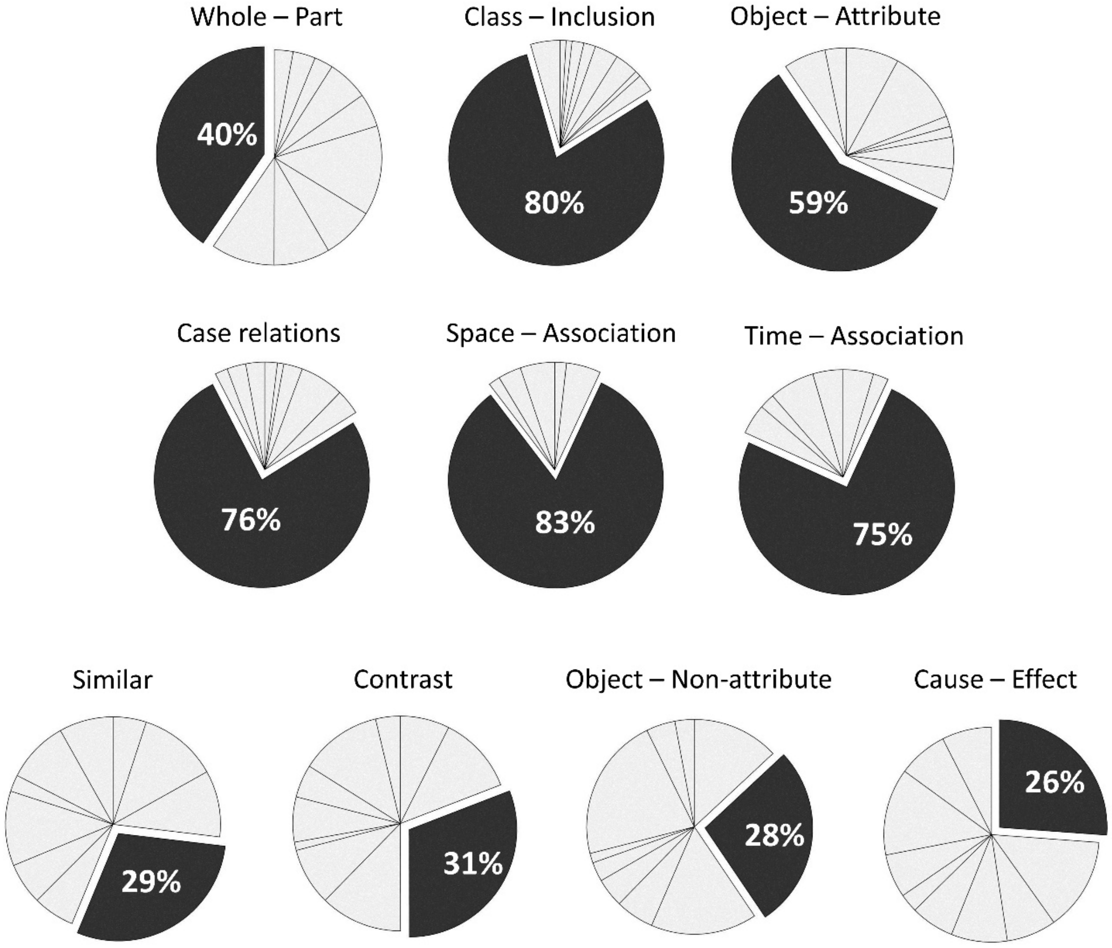


**Supplementary Figure 7. Top-1 accuracy of classifying every word-pair into 1 out of 10 classes of semantic relations.** For every word pair addressed in this study, we calculate the percentage by which it is classified into the correct class of semantic relation (dark gray) vs. an incorrect class of relation (light gray). In the pie chart, different classes of semantic relation are shown as different sectors. In a counterclockwise order, they are whole-part, class-inclusion, object-attribute, case relations, space-association, time-association, similar, contrast, object-non-attribute, cause-effect.

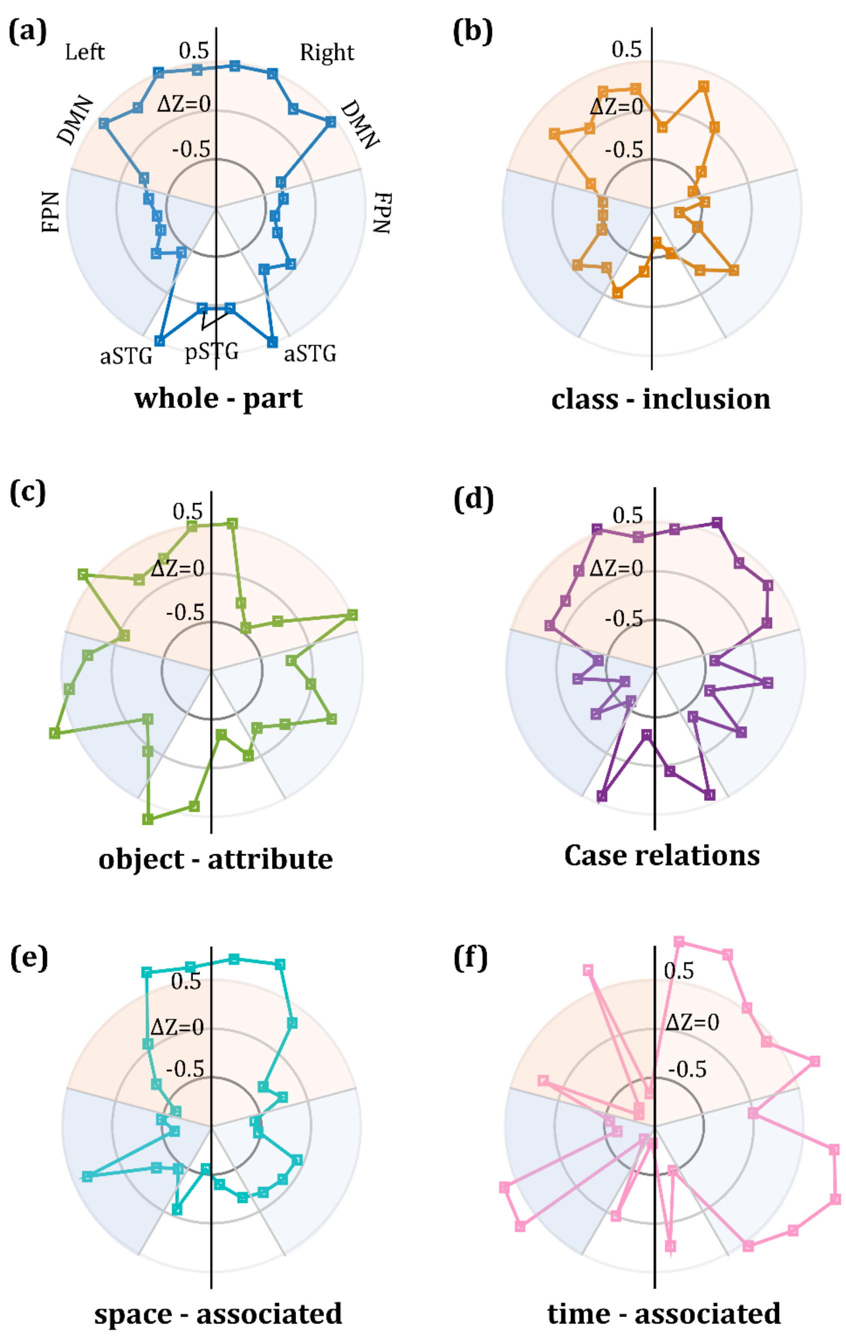


**Supplementary Figure 8. Representational geometry of each semantic relation.** For a given semantic relation, nodes correspond to predefined ROIs as labeled in panel (a); the same labels also apply to panel (b) through (f). The position of each node relative to the center indicates the average projection of a semantic relation onto the ROI.

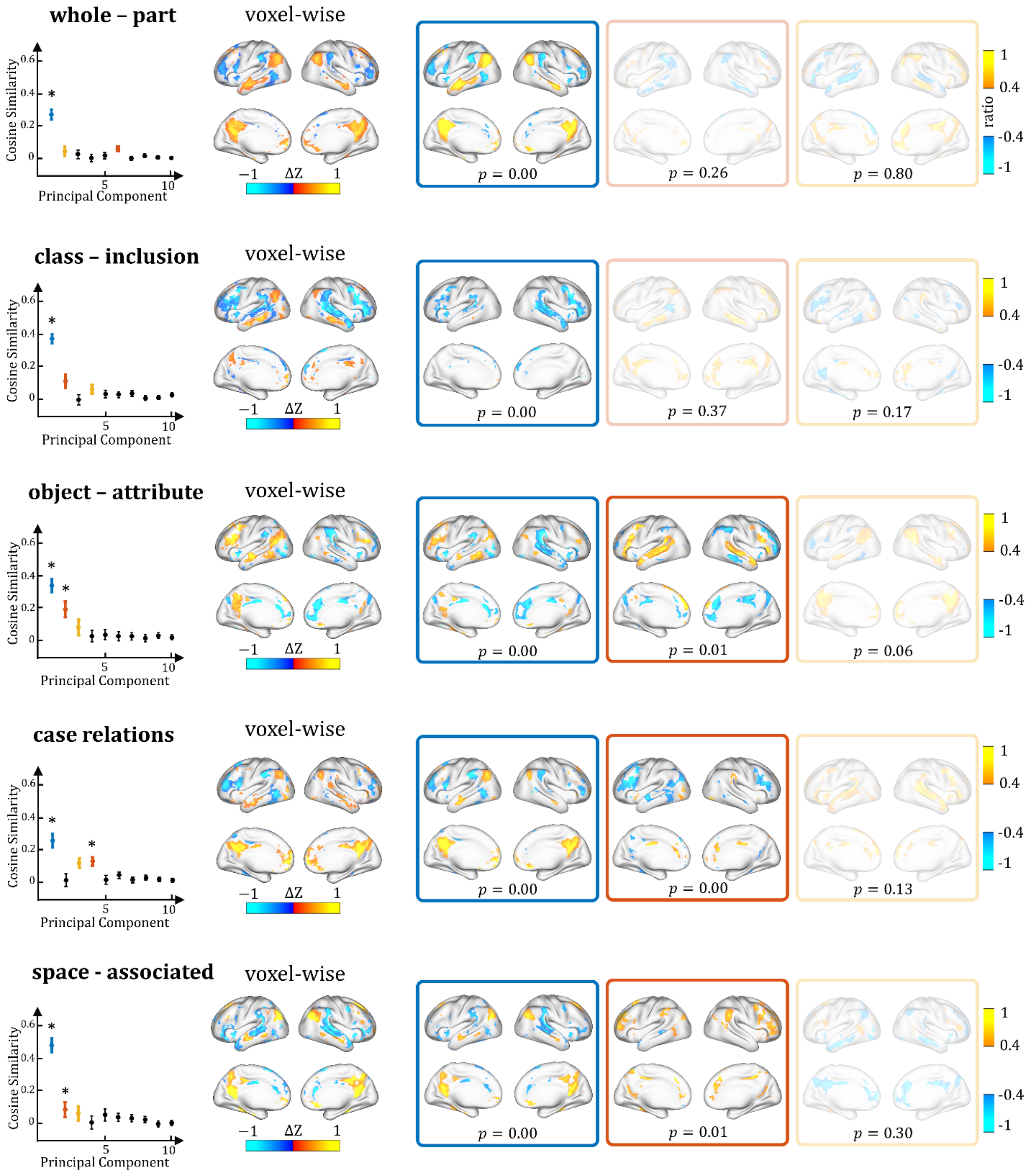

whole – part
class – inclusion
object – attribute
case relations
space - associated
voxel-wise
Cosine Similarity
Principal Component
0
0.2
0.4
0.6
5
10
*
−1
ΔZ
1
ratio
1
0.4
-0.4
-1
p = 0.00
p = 0.26
p = 0.80
p = 0.00
p = 0.37
p = 0.17
p = 0.00
p = 0.01
p = 0.06
p = 0.00
p = 0.00
p = 0.13
p = 0.00
p = 0.01
p = 0.30

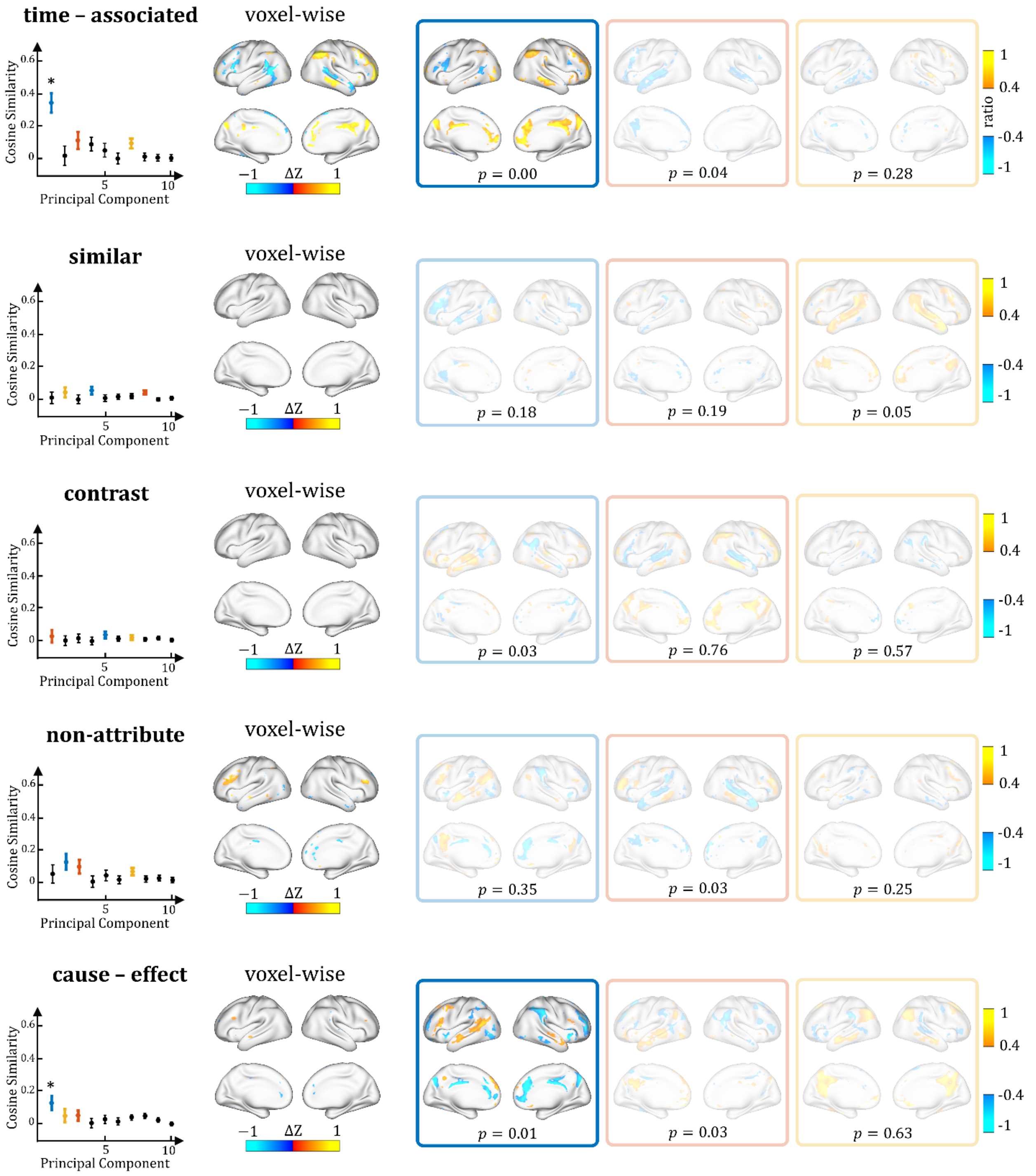


**Supplementary Figure 9. Cortical patterns representing each semantic relation.** Each plot on the left shows the average cosine similarity between the cortical projection of every word pair and each of the top-10 principal components. The error bar indicates the standard error of the mean. The cortical patterns with the top-3 largest average cosine similarities are highlighted in blue, yellow and red in the plots and in the boxes that bound those patterns. The patterns that lack statistical significance (one-sample t-test, two-sided, $p < 0.01$) are shown in faded boxes, whereas the patterns of statistical significance are not faded. In each cortical pattern, the color indicates the voxel value after a min-max normalization to [-1,1]. See details in Supplementary Method 7. For comparison, the cortical map obtained with univariate voxel-wise analysis is shown for each category (paired permutation test, two-sided, FDR $q < 0.05$).

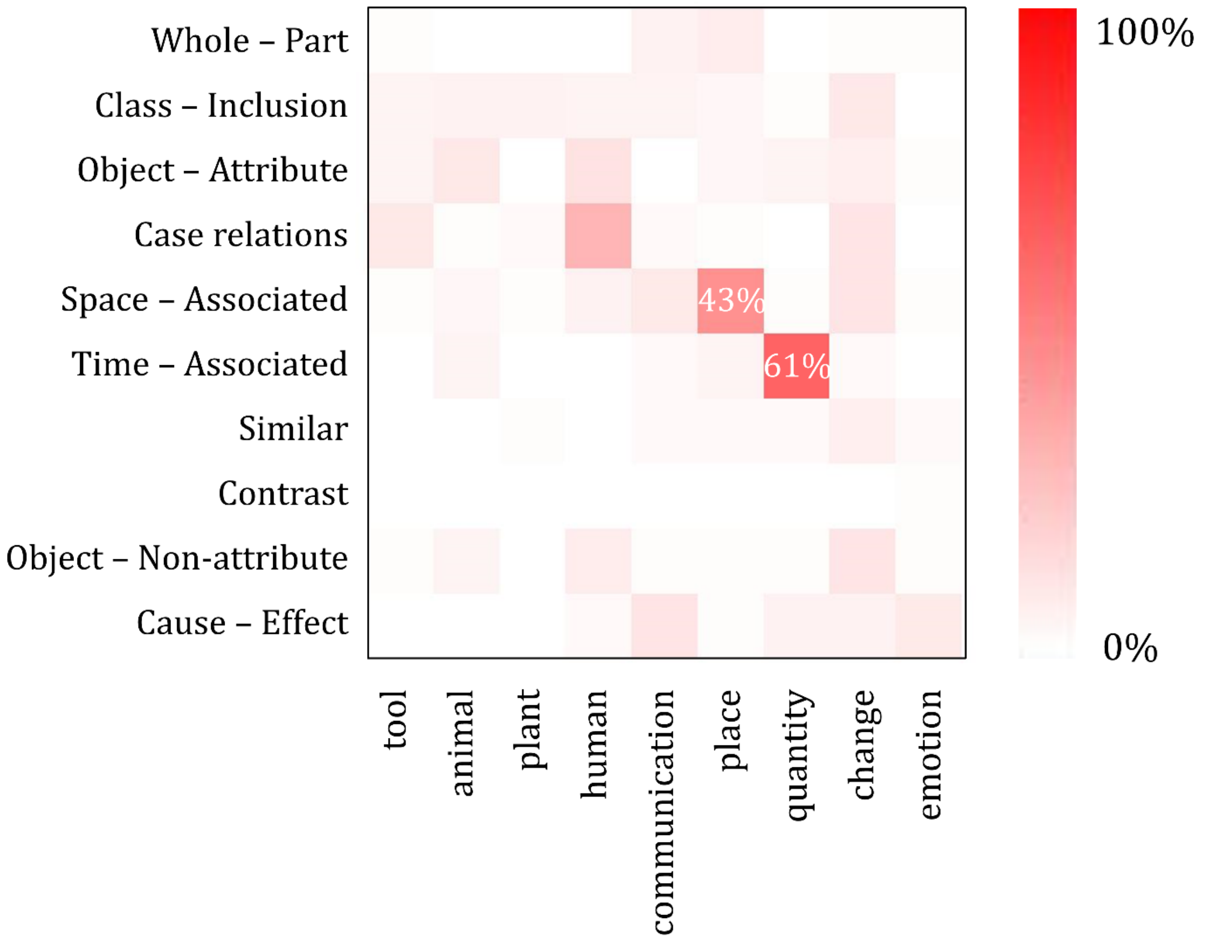


**Supplementary Figure10. The association between semantic relations and semantic categories.** For each relation, we count the number of word-pair samples associated with each category and computed the percentage relative to the total number of word-pairs. The percentage is color coded and displayed in a matrix, where rows correspond to relations and columns correspond to categories. See Supplementary Method 5 for more details about the analysis.

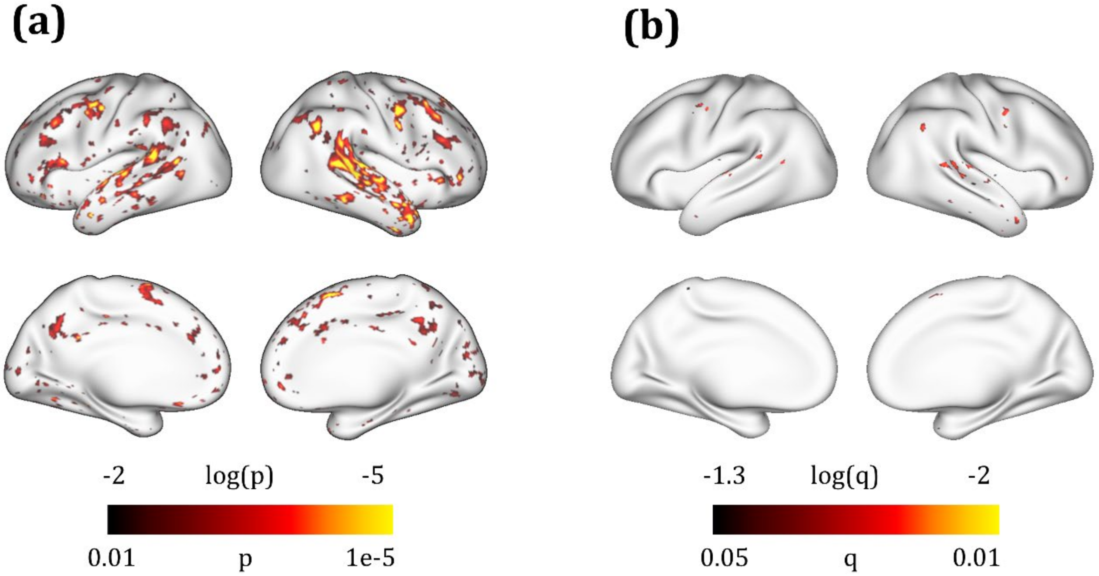


**Supplementary Figure 11. Cortical locations with significant individual variance.** We evaluate the cortical activation with the testing story separately for each subject. The map highlights the voxels where individual variation is significant in a 2-way ANOVA given a threshold with uncorrected $p < 0.01$ (two-sided) or with FDR $q < 0.05$ (two-sided) after correction for multiple comparisons.

### Supplementary Tables

**Supplementary Table 1: Details of frequently appeared words in the training dataset**

| Sub | # words | # unique words | Top 5 appeared words (with word2vec embedding) | | | | |
|---|---|---|---|---|---|---|---|
| 1 | 2,885 | 756 | I (155) | the (113) | my (95) | so (53) | that (51) |
| 2 | 2,416 | 759 | I (100) | the (84) | was (48) | her (48) | that (43) |
| 3 | 2,530 | 813 | I (113) | the (97) | was (84) | my (73) | he (56) |
| 4 | 3,112 | 791 | I (157) | the (141) | in (57) | you (52) | my (50) |
| 5 | 2,745 | 739 | the (140) | I (125) | in (53) | you (52) | my (51) |
| 6 | 3,155 | 867 | the (159) | I (143) | he (77) | was (65) | you (63) |
| 7 | 2,509 | 751 | I (119) | the (114) | like (63) | she (52) | you (52) |
| 8 | 2,574 | 848 | I (150) | the (149) | was (54) | that (51) | you (51) |
| 9 | 2,154 | 623 | I (144) | was (67) | my (63) | the (55) | you (44) |
| 10 | 2,272 | 843 | the (107) | I (81) | that (50) | in (37) | you (29) |
| 11 | 1,699 | 613 | I (94) | the (64) | was (51) | that (41) | in (40) |
| 12 | 2,253 | 765 | the (119) | I (114) | was (60) | that (40) | in (38) |
| 13 | 2,246 | 651 | the (119) | I (117) | you (49) | in (41) | me (41) |
| 14 | 2,976 | 833 | I (155) | the (153) | my (96) | that (73) | she (57) |
| 15 | 2,427 | 764 | I (152) | the (110) | that (67) | was (53) | in (52) |
| 16 | 3,089 | 803 | I (176) | the (111) | that (84) | was (81) | my (59) |
| 17 | 1,966 | 651 | I (124) | the (75) | you (50) | was (46) | my (43) |
| 18 | 2,227 | 704 | I (144) | the (81) | you (47) | was (45) | my (44) |
| 19 | 2,863 | 906 | I (138) | the (136) | my (60) | was (58) | that (51) |

#### Supplementary Table 2: Details of ROI in Figure 3

| Left | | | | | Right | | | | |
|---|---|---|---|---|---|---|---|---|---|
| Name | Location | # voxels | Voxel Corr | ROI Corr | Name | Location | # voxels | Voxel Corr | ROI Corr |
| IFS | −47, 32, 14 | 155 | 0.48 ± 0.11 | 0.58 | IFS | 48, 35, 13 | 117 | 0.34 ± 0.05 | 0.38 |
| SMG | −53, −31, 23 | 530 | 0.29 ± 0.17 | 0.49 | SMG | 55, −26, 26 | 980 | 0.34 ± 0.09 | 0.50 |
| AG | −47, −65, 26 | 808 | 0.33 ± 0.10 | 0.31 | AG | 53, −54, 25 | 560 | 0.42 ± 0.06 | 0.47 |
| STG | −58, −19, −4 | 859 | 0.29 ± 0.08 | 0.24 | STG | 60, −16, −3 | 863 | 0.28 ± 0.10 | 0.24 |
| MT | −38, −81, −5 | 661 | 0.27 ± 0.12 | 0.40 | MT | 40, −78, −7 | 536 | 0.27 ± 0.10 | 0.36 |
| FuG | −38, −34, −25 | 695 | 0.34 ± 0.14 | 0.47 | FuG | 38, −32, −27 | 698 | 0.13 ± 0.14 | 0.11 |
| PhG | −28, −32, −18 | 110 | 0.40 ± 0.12 | 0.51 | PhG | 30, −30, −18 | 118 | 0.26 ± 0.11 | 0.35 |
| PCC | −6, −55, 34 | 377 | 0.43 ± 0.07 | 0.48 | PCC | 6, −54, 35 | 400 | 0.44 ± 0.09 | 0.50 |

#### Supplementary Table 3: Details of each semantic category

| Semantic category | Wordnet synsets | Semantic definition | # sample words | Example words |
|---|---|---|---|---|
| tool | 'tool.n.01' | an implement used in the practice of a vocation | 200 | axe, drill, fork, saw, wrench |
| animal | 'animal.n.01' | a living organism characterized by voluntary movement | 734 | ant, cat, dog, fox, hen, owl |
| plant | 'plant.n.02' | a living organism lacking the power of locomotion | 387 | aloe, beet, corn, lotus, rosewood |
| human | 'adult.n.01'; 'worker.n.01' | a fully developed person; a person who works at a specific occupation | 808 | barber, clerk, groom, hunter, man |
| communication | 'communication.n.02' | something that is communicated by or to or between people or groups | 2,027 | chat, discussion, gossip, idea, speaking |
| place | 'location.n.01'; 'building.n.01' | a point or extent in space; a structure that has a roof and walls and stands more or less permanently in one place | 814 | arena, city, grave, inside, terminal |
| quantity | 'definite_quantity.n.01'; 'measure.n.02' | a specific measure of amount; how much/many of something that you can quantify | 958 | billion, decade, single, gallon, megabyte |
| change | 'change.v.01'; 'change.v.02' | cause to change, make difference; undergo a change | 3,417 | abort, fall, heal, reduce, thicken |
| emotion | 'feeling.n.01' | the experiencing of affective and emotional states | 504 | anxiety, concern, daze, dream, mood |

**Supplementary Table 4: Details of each semantic relation**

| Semantic relationship | Number of paired samples | Number of significant voxels | Example word pairs |
|---|---|---|---|
| whole-part | 178 | 10,607 | hand-finger, zoo-animal, hour-second |
| class-inclusion | 113 | 10,768 | color-green, weapon-spear, tree-oak |
| object-attribute | 63 | 9,037 | fire-hot, child-innocent, heart-beat |
| case relations | 106 | 9,550 | coach-player, writer-story, barber-scissors |
| space-associated | 58 | 11,496 | library-book, mine-coal, mall-shopping |
| time-associated | 44 | 6,129 | morning-sunrise, winter-snow, christmas-presents |
| similar | 160 | 0 | house-home, kid-child, teach-instruct |
| contrast | 162 | 0 | hot-cold, rich-poor, top-bottom |
| object-nonattribute | 69 | 1,124 | fire-cold, slavery-freedom, optimist-despair |
| cause-effect | 107 | 244 | loss-grief, heat-sweat, study-learn |

### Supplementary Reference


1 Maaten, L. v. d. & Hinton, G. Visualizing data using t-SNE. *Journal of machine learning research* **9**, 2579-2605 (2008).

2 Gotts, S. J. *et al.* Two distinct forms of functional lateralization in the human brain. *Proceedings of the National Academy of Sciences* **110**, E3435-E3444 (2013).

3 Knecht, S. *et al.* Language lateralization in healthy right-handers. *Brain* **123**, 74-81 (2000).

4 Hasson, U., Malach, R. & Heeger, D. J. Reliability of cortical activity during natural stimulation. *Trends in cognitive sciences* **14**, 40-48 (2010).

5 Brysbaert, M., Warriner, A. B. & Kuperman, V. Concreteness ratings for 40 thousand generally known English word lemmas. *Behavior research methods* **46**, 904-911 (2014).

6 Miller, G. *WordNet: An electronic lexical database*. (MIT press, 1998).

7 Jurgens, D. A., Turney, P. D., Mohammad, S. M. & Holyoak, K. J. in *Proceedings of the First Joint Conference on Lexical and Computational Semantics-Volume 1: Proceedings of the main conference and the shared task, and Volume 2: Proceedings of the Sixth International Workshop on Semantic Evaluation.* 356-364 (Association for Computational Linguistics).

8 Mikolov, T., Yih, W.-t. & Zweig, G. in *Proceedings of the 2013 Conference of the North American Chapter of the Association for Computational Linguistics: Human Language Technologies.* 746-751.

9 Haxby, J. V. Multivariate pattern analysis of fMRI: the early beginnings. *Neuroimage* **62**, 852-855 (2012).

10 Mikolov, T., Sutskever, I., Chen, K., Corrado, G. S. & Dean, J. in *Advances in neural information processing systems.* 3111-3119.
